# Corticofugal gated recurrency captures auditory cortical responses

**DOI:** 10.64898/2025.12.09.693169

**Authors:** Lorenzo Mazzaschi, Andrew J King, Ben DB Willmore, Nicol S Harper

**Affiliations:** Department of Physiology, Anatomy and Genetics, University of Oxford, Sherrington Building, OX1 3PT, Oxford, United Kingdom

## Abstract

Memory is essential for the neural processing of natural sounds. It has been proposed that cortical memory is subserved by gated recurrency, a powerful machine learning method that enables memory of sequential dependencies. However, standard forms lack biological realism. We built a computational model using a subtractive form of gated recurrency, consistent with cortical circuitry and feedback that resembles corticofugal projections. This architecture outperformed state-of-the-art models with delay lines or non-biological gated recurrency at predicting auditory cortical responses to natural sounds. Its performance was further improved by incorporating a network layer that mirrors the high neuronal density in superficial cortical layers. Analyzing the model, we found that memory retention is longer in secondary than in primary auditory cortex, and that gated recurrency particularly helps capture responses during abrupt changes and quiet periods in the sounds. This work lays out the fundamental functional circuitry of the auditory pathway for processing the sequential dependencies in natural sounds.

## Introduction

A crucial component of neural circuits that underlie sound processing is memory of the sequential dependencies in sounds. Natural sounds in particular contain long sequential dependencies ^1,2^, for example, vowels commonly follow consonants in speech, or footsteps repeat with silence between them. Indeed, neurons in the auditory cortex are sensitive to the temporal context of sounds on timescales of milliseconds to minutes ^3–5^. While anatomy can reveal the broad architecture of such circuits, modelling the physiology is required to determine which circuit features are involved during their natural operation. Hence, a key step towards explaining the functional organization of the auditory pathway is to capture the responses of neurons to natural sounds using anatomically grounded models with memory for natural sound sequences.

A possible biological mechanism for memory is gated recurrency, which involves cyclically interconnected computational units that selectively retain only relevant information by use of gates, and has until recently been the state-of-the-art in machine learning for processing sequential stimuli ^6^. Indeed, standard gated recurrent models have been shown to improve encoding models’ predictions of auditory cortical responses to natural sounds^7^, though these involve biologically unrealistic multiplicative circuitry. However, gated recurrent circuits that use subtractive mechanisms, consistent with inhibitory interneurons locally gating the inputs and outputs of the pyramidal neurons that drive them, have been hypothesized in the cortex^8^. Furthermore, the auditory pathway as a whole is highly recurrent, with descending corticofugal projections extending as far back as the brainstem ^9,10^. We therefore combined a biologically-based model of subtractive gated recurrency with feedback to brainstem-like input to investigate whether this could explain auditory cortical responses to natural sounds and hence elucidate the circuit properties likely to play essential roles in auditory processing.

Existing models of auditory neural responses vary in complexity, from simple linear filters of sound to convolutional neural networks with multiple hidden layers^11–16^. To date, convolutional models achieve the best performance in predicting cortical responses to natural sounds in awake ferrets ^16^. A common aspect of these models is that they have feedforward architectures, relying on delay lines to retain past information ^14,16^. However, there is no biological evidence for delay lines longer than a few tens of milliseconds ^15^, which is insufficient to capture neural memory or sequential dependencies in natural sounds. Recurrent neural network models are a potential solution to this problem, with gated recurrent networks being a particularly powerful form.

To evaluate memory mechanisms in models of auditory cortical neurons, we fitted a range of neural network architectures (including convolutional, recurrent, and gated recurrent) to neural activity recorded in ferret primary and secondary auditory cortex in response to natural sounds. Amongst the gated recurrent models, some used established architectures from the machine learning domain, while others were directly inspired from the brain, allowing us to ascertain the importance of biological features and known anatomical circuitry.

In particular, we developed a new subtractive gated recurrent network architecture that mimics corticofugal projections to previous stages of the auditory pathway. This model has a gate operating directly on its input, controlled by feedback connections, and provided the best prediction performance across three datasets of ferret auditory cortex responses to natural sounds, outcompeting state-of-the-art models. Its performance on primary auditory cortex data improved further by adding a feedforward expansion layer inspired by the columnar structure of the cortex. This work demonstrates the role provided by key features of the architecture of the auditory pathway in processing the sequential dependencies in natural sounds.

## Results

We quantified model performance on three publicly available datasets of extracellular spiking activity recorded from the auditory cortex of awake, passively listening ferrets presented with wide-ranging libraries of natural sounds. Two sets of responses were recorded to the same ensemble of 591 1-second long stimuli; NAT4-A1 (383 units) was recorded in primary auditory cortex (A1), and NAT4-PEG (195 units) in the posterior ectosylvian gyrus (PEG, a secondary auditory area) ^16^. The third set (NS2, 180 units) was recorded in A1 in response to 299 4-second long stimuli ^17^.

We fitted models to predict neural responses (peri-stimulus time histograms, PSTHs, of the recorded spike trains) from cochleagrams (Supplementary Fig. S1), spectrotemporal representations of the sound stimuli that reflect cochlear filtering properties ^18^. The PSTHs and cochleagrams were binned at 4 ms. Throughout, the term ”stimulus-response snippet” refers to a stimulus and its associated response.

We developed five gated recurrent architectures (Supplementary Figs. S2 and S3; Methods). Gates are sigmoidal functions, which operate by multiplication on memory and incoming input to regulate the flow of information across time steps, determining the temporal integration properties of the recurrent units. The two most widely used gated units are the long short-term memory (LSTM) unit^19^ and the gated recurrent unit (GRU) ^20^, which use 3 and 2 gates, respectively. We developed models using both of these, as well as the minimal gated unit (MGU, containing one gate) ^21^ and the subtractive LSTM (subLSTM) unit^8^. The subLSTM unit, an adaptation of the LSTM unit whose gates operate in a more biologically realistic manner (subtractively rather than multiplicatively), is of particular interest - its architecture was designed to match a potential neural circuit across the layers of the cortex^8^.

We developed a new architecture, the corticofugal LSTM (f-LSTM) model, which uses subLSTM units with feedback connections to an additional gate at the model’s input (Fig. 1a). Corticofugal projections from the auditory cortex are found at nearly all subcortical stages of the auditory pathway, including the medial geniculate body (MGB) in the thalamus, the inferior colliculus (IC) in the midbrain, and the cochlear nucleus (CN) in the brainstem ^9,10^ (Fig. 1b). They have been shown to influence sound processing and learning ^22–26^. The f-LSTM network was designed to mimic these corticofugal components. The stimulus cochleagram is fed to a layer of subLSTM units via a set of delay lines. The subLSTM units project forward to the output layer providing PSTH predictions, but also feed into an *input gating* mechanism operating multiplicatively on the stimulus directly. There is some evidence in mice that corticofugal projections can have a multiplicative/divisive impact on their targets^27^.

**Figure 1.**
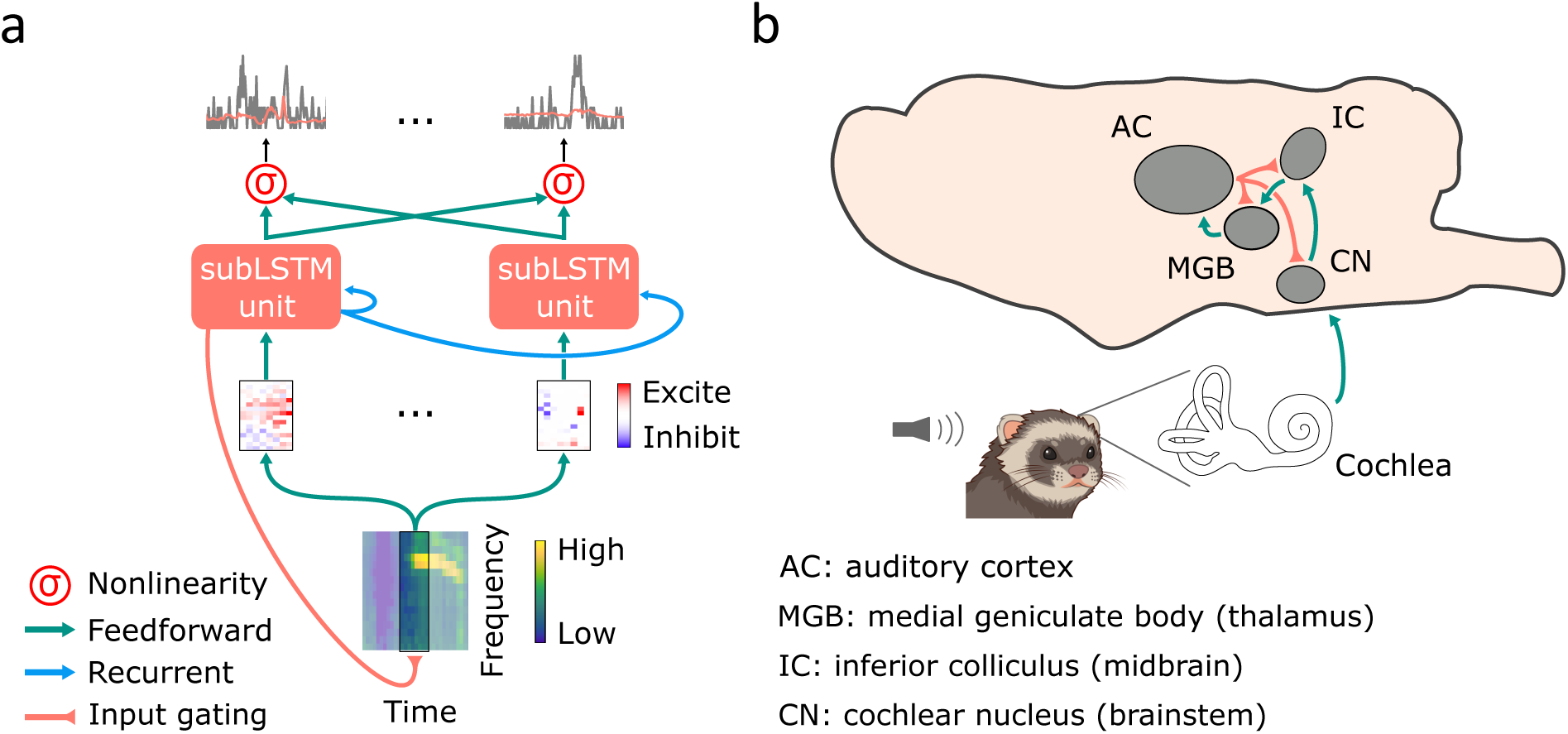
Macrocircuits of the auditory pathway and a model inspired by them. (a) Schematic of the corticofugal long short-term memory (f-LSTM) model architecture. A sound cochleagram is fed into the model at the input (bottom), and is processed by a gated recurrent layer of subtractive LSTM (subLSTM) units, which are inspired by cortical architecture^8^. The input to this gated recurrent layer is the linear mapping of recent cochleagram activity. The prediction of activity for each recorded unit was estimated from the subLSTM units’ activations by a linear mapping followed by a static nonlinearity. The feedback from the gated recurrent layer to the input gating mechanism simulates the effect of corticofugal projections. (b) Schematic of the ferret auditory pathway, showing major processing stations and their associated projections.

The convolutional models of Pennington and David were fitted to whole populations of recorded units at once, as opposed to single-neuron architectures, which are only trained on data from one unit. This enabled model units to learn shared representations in spectrotemporal space, improving model performance ^16^. We therefore implemented all of our architectures as population models.

We also implemented several feedforward models against which to test the benefits of gated recurrency (Supplementary Fig. S2; Methods). A population linear-nonlinear (LN) model provided a simple, well-established architecture as a baseline for comparison^12,16^. In addition, we used the network receptive field (NRF) model from Harper et al. (2016), implemented as a population architecture, as well as three convolutional models from Pennington and David (2023), to provide leading feedforward models to compare against. We followed the nomenclature from Pennington and David (2023): the 1D-CNN (1-Dimensional CNN) model is a neural network with one layer of 1-dimensional convolutional filters; the 1D-x2-CNN model has two hidden layers of 1D convolutional filters; the 2D-CNN (2-Dimensional CNN) model has 3 hidden layers with 2-dimensional convolutional filters. Finally, we also used a network with basic recurrent units (RNN model), which contain no gates ^28^.

### The f-LSTM model is the best predictor of A1 activity

We fitted the models to the stimulus-response relationships of recorded auditory cortical units and then tested their ability to predict responses to held-out stimuli not used in model fitting. For this testing, we used a subset of stimuli that were presented to the animals over several repetitions, to factor in the reliability of each unit’s response. Example units show how different patterns of neural activity are captured differently by the models (Figs. 2a,b). One of the units responded mainly to high-intensity components within the sound, and was well predicted by the LN model (Fig. 2a). The other was active during quieter periods in the sound, and was much better predicted by the f-LSTM than the LN model (Fig. 2b).

**Figure 2.**
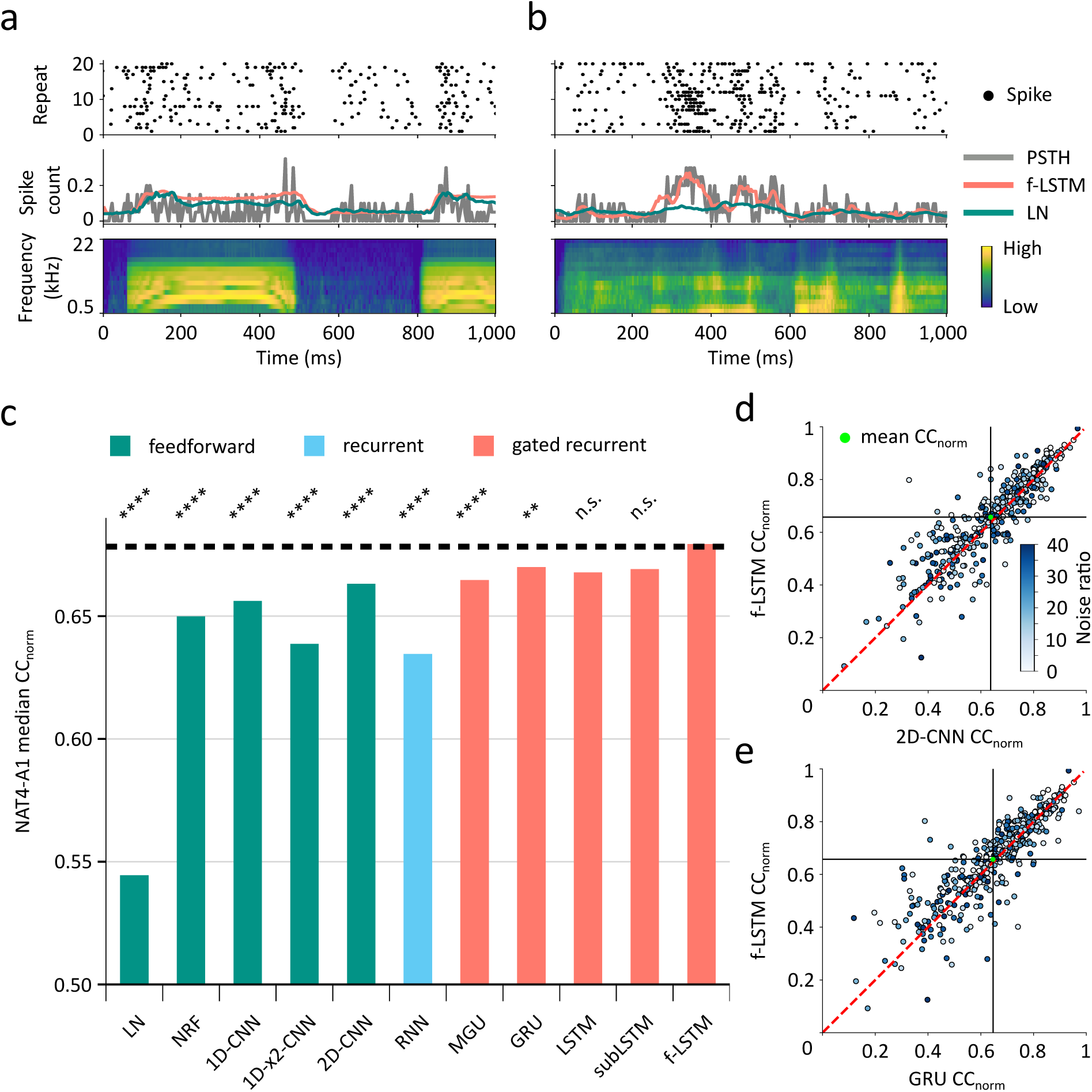
Gated recurrent models provide the best predictions of neural activity in the NAT4-A1 dataset. (a, b) Two example units from the NAT4-A1 dataset. Each panel shows a cochlea-gram of the stimulus played to the ferrets (bottom), the unit’s response overlaid with the overall worst (LN) and best (f-LSTM) model’s prediction (middle), and the raster plot of spikes recorded in response to each repeat of the stimulus (top, from which the PSTH in the middle panel is computed). The unit in (a) is well predicted by both the LN and f-LSTM models, whereas the unit in (b) is well-predicted by the f-LSTM model only. The identifiers for the units are found in the Supplementary Information. (c) Median prediction performance across the dataset for all models, color-coded by model type. The f-LSTM is the best model overall, and all gated recurrent models perform better than all other models. Asterisks indicate a significant difference (Wilcoxon signed-rank test, n=383; see Methods) between each of the models and the f-LSTM. (d) Unit-by-unit comparison of the f-LSTM model with the best feedforward model (2D-CNN). (e) Comparison of the f-LSTM model with the second-best model overall (GRU). Each dot in the scatter plots corresponds to a recorded unit, with its CC_norm_ values for the two models plotted against one another. Units are color-coded by their noise ratio (see Methods). Solid black lines and green dots show mean population performance. Dashed red lines indicate equal performance. The plots show that the majority of units are better predicted by the f-LSTM than both the 2D-CNN and the GRU models.

We evaluated model prediction performance with the normalized correlation coefficient (CC_norm_) between actual and predicted neural activity^29,30^ (see Methods). This measure uses responses to several repeats of the same stimuli to assess the reliability of each recorded unit, normalizing the raw correlation coefficient (CC_raw_) by the unit’s maximum explainable activity. We computed the CC_norm_ for each unit, and summarized model performance by the median CC_norm_ across the population (Fig. 2c for the NAT4-A1 dataset).

The f-LSTM model is the best predictor of neural activity for the NAT4-A1 dataset, achieving a median CC_norm_ of 0.68 and significantly surpassing all but two (LSTM, subLSTM) of the other models (Fig. 2c). Crucially, it significantly improves upon the best feedforward model, the 2D-CNN. All gated recurrent models perform better than all other models, with all comparisons being significant except between the MGU and 2D-CNN models. Convolutional models still reach good levels of performance, confirming the findings of Pennington and David (2023). The NRF model, whose architecture is very similar to the CNN-based ones, performs similarly to the latter on average. The LN model, lacking in computational complexity, is substantially worse than all other models. Finally, the RNN model is the second worst, highlighting the importance of the long memory conferred by gates.

For all the models, we explored different spans of delay lines on the input, picking the span with the highest performance on the validation set, a subset of data held out from training to evaluate models in the development phase. For the gated recurrent models, this resulted in a span of 32 ms, which is close to experimentally observed peak latencies in ferret auditory cortex ^31^; the recurrent model and the feedforward models required longer spans, up to 250 ms, as in previous publications ^14,16^. Therefore, the gated recurrent models perform better than feedforward ones using much shorter and biologically realistic spans. For each model, we used the same delay line span across all three datasets.

While the median CC_norm_ provides a good summary of model performance, it is important to compare models across units to see if the differences are consistent (Figs. 2d,e). For the majority of recorded units, the f-LSTM model produces higher CC_norm_ values than both the best feedforward model (2D-CNN) and the second-best model overall (GRU). Specifically, it surpasses the 2D-CNN model in 66% (n = 383) of the units, and the GRU model in 56% of the units. This is evidence that improvements due to gated recurrency are well distributed over the neural population.

A similar picture was found for the NS2 dataset (Fig. 3), again illustrated for two example units (Figs. 3a,b) and by a comparison of the overall performance of different models (Figs. 3c-e). The f-LSTM once again emerges as the best model overall (0.67 median CC_norm_), with a significant improvement compared to the second-best model (MGU, Fig. 3c). The difference between all gated recurrent models and all feedforward ones is larger in the NS2 than in the NAT4-A1 dataset (the only non-significant comparison is between the GRU and 2D-CNN models), but all other aspects of the results are similar. The improvement due to the f-LSTM model is distributed across the neural population for the NS2 dataset as well (Figs. 3d,e). The f-LSTM model surpasses the best feedforward model (2D-CNN) for 66% (n = 180) of the recorded units, and the best other model (MGU) for 61% of the units.

**Figure 3.**
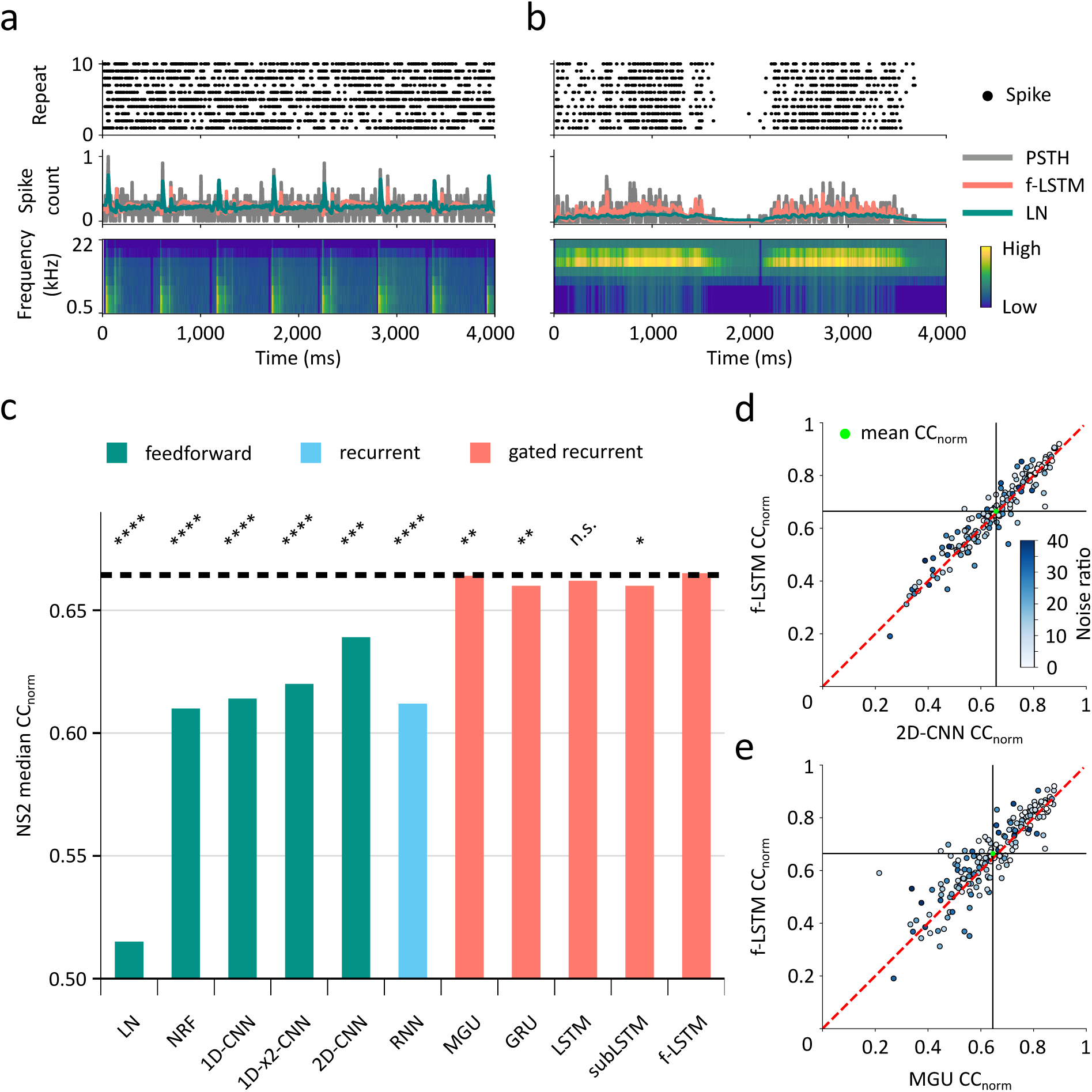
Gated recurrent models provide the best predictions of neural activity in the NS2 dataset. Panels as in Fig. 2. (a, b) Two examples of neural activity from the NS2 dataset. The unit in (a) is well predicted by both the LN and f-LSTM models, whereas the unit in (b) is well-predicted by the f-LSTM model only. The identifiers for the units are found in the Supplementary Information. (c) Median prediction performance by model type. As in the NAT4-A1 dataset, the f-LSTM is the best model overall, and all gated recurrent models perform better than all other models. Asterisks indicate a significant difference (Wilcoxon signed-rank test, n=180) between each of the models and the f-LSTM. (d) Unit-by-unit comparison of the f-LSTM model with the best feedforward model (2D-CNN). (e) Comparison of the f-LSTM model with the second-best model overall (MGU). The plots show that the majority of recorded units are better predicted by the f-LSTM than both the 2D-CNN and the MGU models.

### Cortical depth and gated recurrency

The structure of subLSTM units is inspired by the columnar architecture of the neocortex, with their memory cells corresponding to recurrently connected pyramidal neurons in cortical layer 5 (L5) ^8^. L5 inputs (from layers 2/3 (L2/3) and layer 4 (L4)) and outputs are gated by subtractive, basket cell-like inhibition^8^. In addition, there is considerable evidence that corticofugal projections to MGB and IC originate within L5 and layer 6 (L6) ^23,32,33^. Neurons in L4 and L2/3, on the other hand, are the primary recipients of afferent input and project primarily within the cortex, including to L5 and L6^32,33^. Given that the architecture of the f-LSTM model was inspired by corticofugal projections, we hypothesized that deep neurons within cortex would be predicted better by the f-LSTM model than superficial neurons.

We therefore plotted the CC_norm_ of recorded units against their depth, for both the f-LSTM and 2D-CNN models, on the NAT4-A1 and NS2 datasets (Figs. 4a,b). Indeed, the difference in CC_norm_ between the f-LSTM and the 2D-CNN models is greater for deeper units, reaching its maximum between 200 µm and 400 µm below the L3/L4 border for both datasets. Depth values were available for only a subset of units from each dataset, covering similar ranges for NAT4-A1 and NS2 (Fig. 4c). Each unit was assigned to a laminar group depending on its depth: either L2/3, L4, or L5/6^16^. The difference between the f-LSTM and 2D-CNN models’ performance is maximal for units recorded in L5/6.

**Figure 4.**
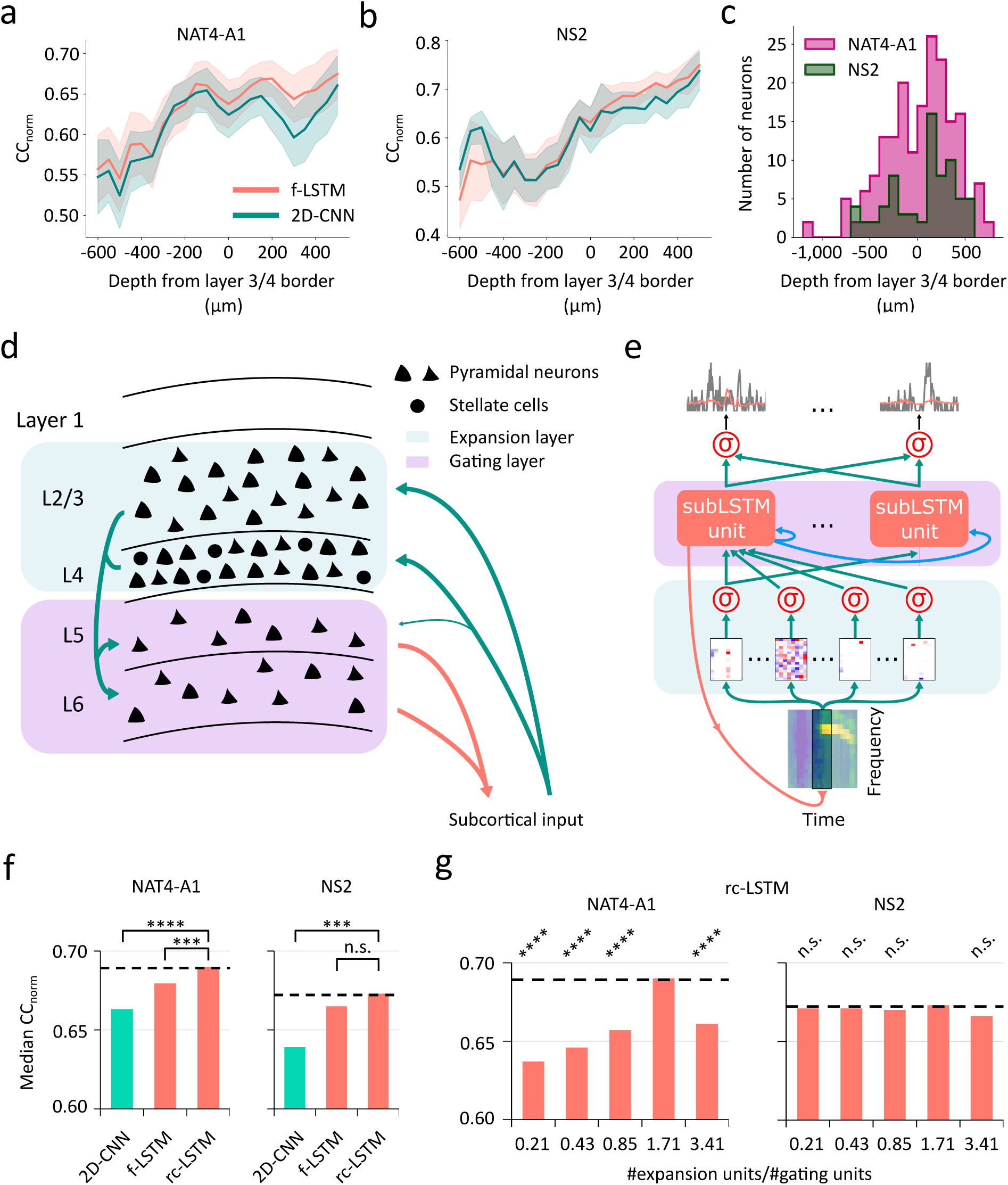
Relative model performance depends on depth of neurons in auditory cortex, and incorporating biological detail about cortical layers in the f-LSTM model improves performance. (a, b) Model performance (CC_norm_) plotted against recorded unit depth for the f-LSTM and 2D-CNN models, for the NAT4-A1 and NS2 datasets, respectively. The difference between the two models is larger for deep (L5/6) units in both datasets. In (a), the difference between the f-LSTM and the 2D-CNN model is significantly higher for L5/6 than for L2/3 and L4 units (unpaired t-test, n1=36, n2=142, t=2.03, df=176, p=0.04). In (b), there are no significant differences (unpaired t-test, n1=20, n2=52, t=0.42, df=70, p=0.68). CC_norm_ values were averaged over 200 µm-wide depth windows overlapping by 50 µm. The shaded areas show the standard error of the mean. (c) Distribution of recorded units across the range of depths for the NAT4-A1 and NS2 datasets. The distributions are overlapping except for 15 units in NAT4-A1. For comparability, we excluded these from the plot in (a). For the results using the full range of depths in NAT4-A1, as well as the results for NAT4-PEG, see Supplementary Fig. S4. (d) Schematic of a cortical column showing microcircuits and high neuronal density in L2/3 and L4 compared to L5 and L6; L2/3 and L4 act as a dimensionality expansion mechanism. (e) Schematic of the rc-LSTM model showing potential correspondence with the cortical column via shading of network layers; a feedforward expansion layer was added to the f-LSTM model’s structure to mirror dimensionality expansion. (f) rc-LSTM model prediction performance on NAT4-A1 and NS2; the rc-LSTM model surpasses the f-LSTM and 2D-CNN models on both datasets. Asterisks indicate a significant difference (Wilcoxon signed-rank test, n=383 for NAT4-A1, n=180 for NS2). (g) rc-LSTM model performance when varying the number of units in the expansion layer, from 32 (21% of the 150 units in the gating layer, dimensionality reduction) to 512 (341%, super-expansion); on both datasets, the best performance is given by a layer with 256 units, denoting an expansion. Asterisks indicate a significant difference (Wilcoxon signed-rank test, n=383 for NAT4-A1, n=180 for NS2) with the 256-unit model.

Encouraged by the correspondence between the f-LSTM model’s performance and neuronal depth, we attempted to incorporate an additional biological element into the network. It has been shown that L2/3 and L4 of the ferret auditory cortex have much higher neuronal densities than L5 and L6^34^ (Fig. 4d). To parallel the connection from high neuronal density L2/3 and L4 to lower density L5 and L6, we modified the f-LSTM model by adding a high-dimensional feedforward layer between the model’s input and the gated recurrent layer. This layer, containing 256 units, provides a dimensional expansion compared to the gated recurrent layer (150 units, see Supplementary Information). We termed this layer the *expansion* layer, and the resulting architecture the ReConfiguration LSTM (rc-LSTM, Fig. 4e) model: the expansion prior to the gated recurrent (*gating* ) layer reconfigures the input into high-dimensional space.

The rc-LSTM model improves upon the f-LSTM model in both datasets (albeit non-significantly in NS2; Fig. 4f). It reaches a median CC_norm_ of 0.69 and 0.67 in NAT4-A1 and NS2, respectively. To verify whether this improvement was really the result of dimensionality expansion, or simply due to the addition of an extra layer in the network, we also varied the number of units in the added layer, from 32 (super-reduction) to 512 (super-expansion). For both datasets, the best result is with 256 units, providing a 71% dimensionality increase compared to the 150-unit subLSTM layer it projects to (Fig. 4g). This matches almost exactly experimental findings in ferrets of the relative neuronal population sizes between superficial and deep layers of auditory cortex ^34^. In NAT4-A1, the differences between the prediction performance of the 256-unit rc-LSTM model and all other rc-LSTM models are significant; in NS2, none of the differences are significant.

### PEG: model performance and temporal integration properties

To test whether gated memory mechanisms are also beneficial in higher-level auditory cortex, we fitted the models to the NAT4-PEG dataset. The activity of two example PEG units responding to sound onset (Fig. 5a) or to short quiet gaps between low-frequency sound components (Fig. 5b), respectively, is captured to different degrees by the f-LSTM and the LN models. The ranking of model performance observed for the A1 datasets also partly applies for NAT4-PEG (Fig. 5c): the f-LSTM model provides the best prediction performance on this dataset (0.59 median CC_norm_). It is significantly better than all other models, with a larger difference from the best feedforward model (1D-CNN) than in either A1 dataset. The subLSTM model also performs well. However, other forms of gated recurrency appear to be less useful: the 1D-CNN model performs (non-significantly) better than the MGU, GRU, and LSTM models. Furthermore, the rc-LSTM model does not improve upon the f-LSTM model for this dataset, and the performance of all models is worse than for the A1 datasets, which is consistent with the results of Pennington and David (2023).

**Figure 5.**
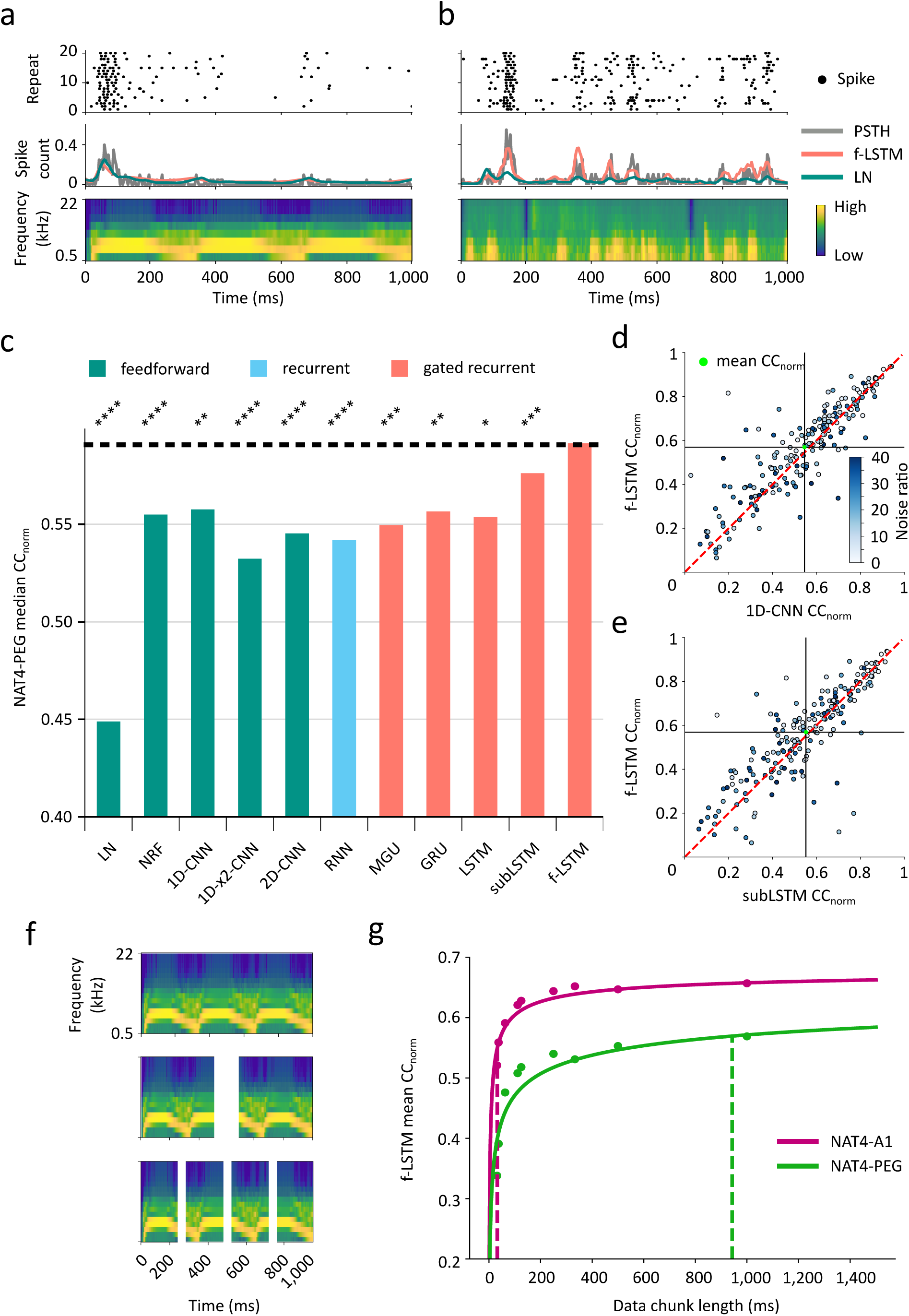
The f-LSTM model is the best predictor of neural activity in PEG, and neurons in this cortical area integrate information on longer timescales than those in A1. (a)-(d) as in Figure 2. (a, b) Two examples of neural responses from the NAT4-PEG dataset. The unit in (a) is well predicted by both the LN and f-LSTM models, whereas the unit in (b) is well-predicted by the f-LSTM model only. The identifiers for the units are found in the Supplementary Information. (c) Median prediction performance by model type. As in the NAT4-A1 dataset, the f-LSTM is the best model overall. Asterisks indicate a significant difference (Wilcoxon signed-rank test, n=195) between each of the models and the f-LSTM. (d) Unit-by-unit comparison of the f-LSTM model with the best convolutional model (1D-CNN). (e) Comparison of the f-LSTM model with the second-best model overall (subLSTM). The plots show that the majority of recorded units are better predicted by the f-LSTM than either of these models. (f) Data ”chunking”: stimulus-response snippets were split into several equal-length chunks, and the f-LSTM model was re-fitted to the NAT4-A1 and NAT4-PEG datasets for each split length. This example shows the 1/2 and 1/4 splits. (g) Mean CC_norm_ for the f-LSTM model plotted against chunk length, for the NAT4-A1 and NAT4-PEG recordings. The solid lines show power-law fits, whereas the dashed vertical lines indicate the chunk length at which the CC_norm_ reaches 80% of its maximum value. This value is more than one order of magnitude larger in PEG than in A1.

For the majority of PEG units, the f-LSTM model performs better in predicting their responses than the 1D-CNN (best feedforward model) and subLSTM (second-best model overall) (Figs. 5d,e). Specifically, it surpasses the 1D-CNN model in 61% (n = 195) of the units, and the subLSTM model in 59% of the units. However, a few PEG units fall very far on either side of the red equal performance line, indicating that they benefit greatly from the use of one architecture over the other. This is not as evident in the NAT4-A1 and NS2 datasets, suggesting that neurons in PEG perform more distinct operations than in A1.

To investigate the use of memory in auditory cortex, we looked at the temporal integration properties of A1 and PEG neurons. Using the NAT4-A1 and NAT4-PEG datasets, we first split each stimulus-response snippet into equal-duration chunks (by 1/2, 1/3, 1/4, 1/8, 1/9, 1/16, 1/27, and 1/32; Fig. 5f). We then fitted the f-LSTM model to the dataset for each chunk length, measuring its performance using the mean CC_norm_ across units (Fig. 5g). We chose the f-LSTM because it is the best model across datasets and has arbitrarily long memory conferred by gated recurrency, meaning it is the length of the data chunk that sets an upper limit on the memory the model can learn to use. For both datasets, prediction performance improves as chunk length increases (Fig. 5g). However, the performance reaches a maximum at longer chunk lengths in PEG than in A1, suggesting that PEG neurons integrate information over longer timescales than A1 neurons. We fitted the increase with a power-law function, which allowed us to calculate a proxy of a time constant for each dataset. The vertical dashed lines indicate the chunk length *L*_0.8_ at which model performance reaches 80% of its maximum, with values of *L*_0.8_ = 32 ms and *L*_0_._8_ = 943 ms for A1 and PEG, respectively. This suggests that PEG operates at timescales much longer than A1. In the NS2 dataset (recorded in A1), with data chunks up to 4 seconds long, we found *L*_0_._8_ = 43 ms (Supplementary Fig. S5).

### Some stimulus features potentially driving performance improvements

Gated memory is clearly beneficial for modelling neuronal responses in auditory cortex. To gain an insight into what might drive performance improvements across the population, we investigated recorded units that show a large increase in CC_norm_ for the consistently best gated model (the f-LSTM) compared to the best feedforward model (the 2D-CNN and 1D-CNN for the NAT4-A1 and NAT4-PEG datasets, respectively). Two activity patterns were often exhibited by the units that showed the greatest improvements in performance in each dataset. First, some units respond strongly to abrupt changes in sound level within the natural sounds - including auditory boundaries, sound onsets, offsets, and similarly sharp changes (Fig. 6a). Second, some of the units are sensitive to the duration of quiet and silent periods within the stimuli, increasing their activity as the period of quiet increased (Fig. 6b).

**Figure 6.**
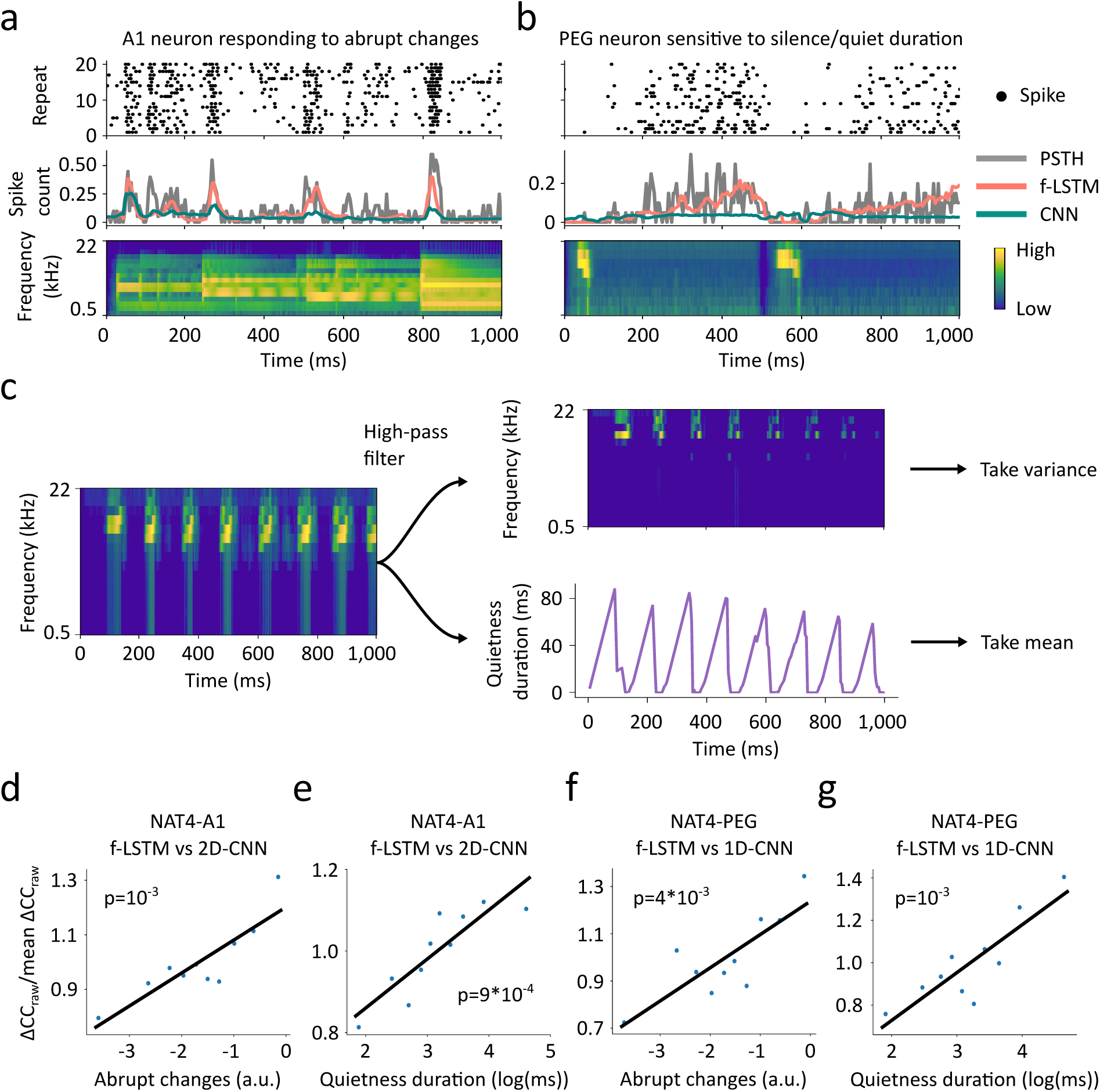
Gated recurrency improves response prediction over feedforward networks particularly for sounds containing long quiet periods and abrupt level changes. (a) Example A1 unit responding to abrupt changes in the stimulus level, which was well captured by the f-LSTM but not by the 2D-CNN model. (b) Example PEG unit responding to the duration of quiet periods in the stimulus, well captured by the f-LSTM but not by the 1D-CNN model. The identifiers for the units in (a) and (b) are found in the Supplementary Information. (c) Calculation of abrupt level changes (top right) and quietness duration (bottom right) measures for an example stimulus (middle left). (d) NAT4-A1 CC_raw_ change plotted against abrupt stimulus level changes. (e) NAT4-A1 CC_raw_ change plotted against stimulus quietness duration. (f) NAT4-PEG CC_raw_ plotted against abrupt stimulus level changes. (g) NAT4-PEG CC_raw_ plotted against stimulus quietness duration. In (d), (e), (f), and (g), data are plotted by decile against the natural log of the respective measures. The solid black line is the line of best fit (ordinary least squares fit) and the p-value (unpaired t-test, n=10, df=8) of its coefficient is reported on the plot. The value of the t-statistic is, respectively: t=5.01 (d), t=5.12 (e), t=3.99 (f), t=4.85 (g).

To quantify abrupt changes and quietness duration in each stimulus, we developed two measures (Fig. 6c; see Methods). We calculated a quietness counter for each stimulus by counting the number of quiet time bins since the last bin of loud sound. The quietness duration is defined as the mean of this quietness counter. For the abrupt changes measure, we computed a high-pass filtered version of the stimulus, whose variance indicates the prevalence of abrupt changes in sound level.

We then attempted to relate these measures to improvements in model performance across the neuronal population. Specifically, we plotted the change in the raw correlation coefficient (CC_raw_) between the f-LSTM and the best feedforward model against the abrupt changes and quietness duration measures, for both the NAT4-A1 and NAT4-PEG datasets (Figs. 6d-g). We used the CC_raw_ for this analysis to be able to use all the stimuli and not just the ones that were played to the ferrets over multiple repeats. Data are plotted by deciles, each including 10% of the stimuli used in the dataset. The CC_raw_ difference was averaged across recorded units for each stimulus and normalized by the mean CC_raw_ difference across all the stimuli and the neuronal population. In all four cases, CC_raw_ difference is significantly correlated with the stimulus measure, with the quietness duration measure exhibiting the strongest and most significant correlations in each dataset (Figs. 6d-g). Overall, these results suggest that part of the improvement observed with the f-LSTM model over the state-of-the-art feedforward ones may emerge from capturing sensitivity to specific stimulus features such as the duration of quiet periods and the presence of abrupt changes in sound level.

### The f-LSTM and rc-LSTM match other gated recurrent models on benchmark tasks

Gated recurrency was first developed to solve problems in machine learning where sequential data processing was required ^35^. Therefore, we evaluated our two new gated architectures on a sequential machine learning task. Sequential tasks, requiring memory, are a common benchmark to test the viability of recurrent and gated recurrent architectures. The well-known MNIST dataset of handwritten digits, used to train models to classify digits independently of their specific handwriting, can be adapted for use in a sequential task^8,36^. 2D images of 28x28 pixels depicting digits 0-9 can be flattened into 1D vectors and fed sequentially to models rather than all at once (Fig. 7a). Using a pre-existing pipeline, we adapted the f-LSTM and the rc-LSTM architectures so that they could be trained for sequential MNIST classification. Overall, our two new models perform comparably well to the LSTM model (Fig. 7b). This was the case for the subLSTM model too, confirming the results from Costa et al. (2017).

**Figure 7.**
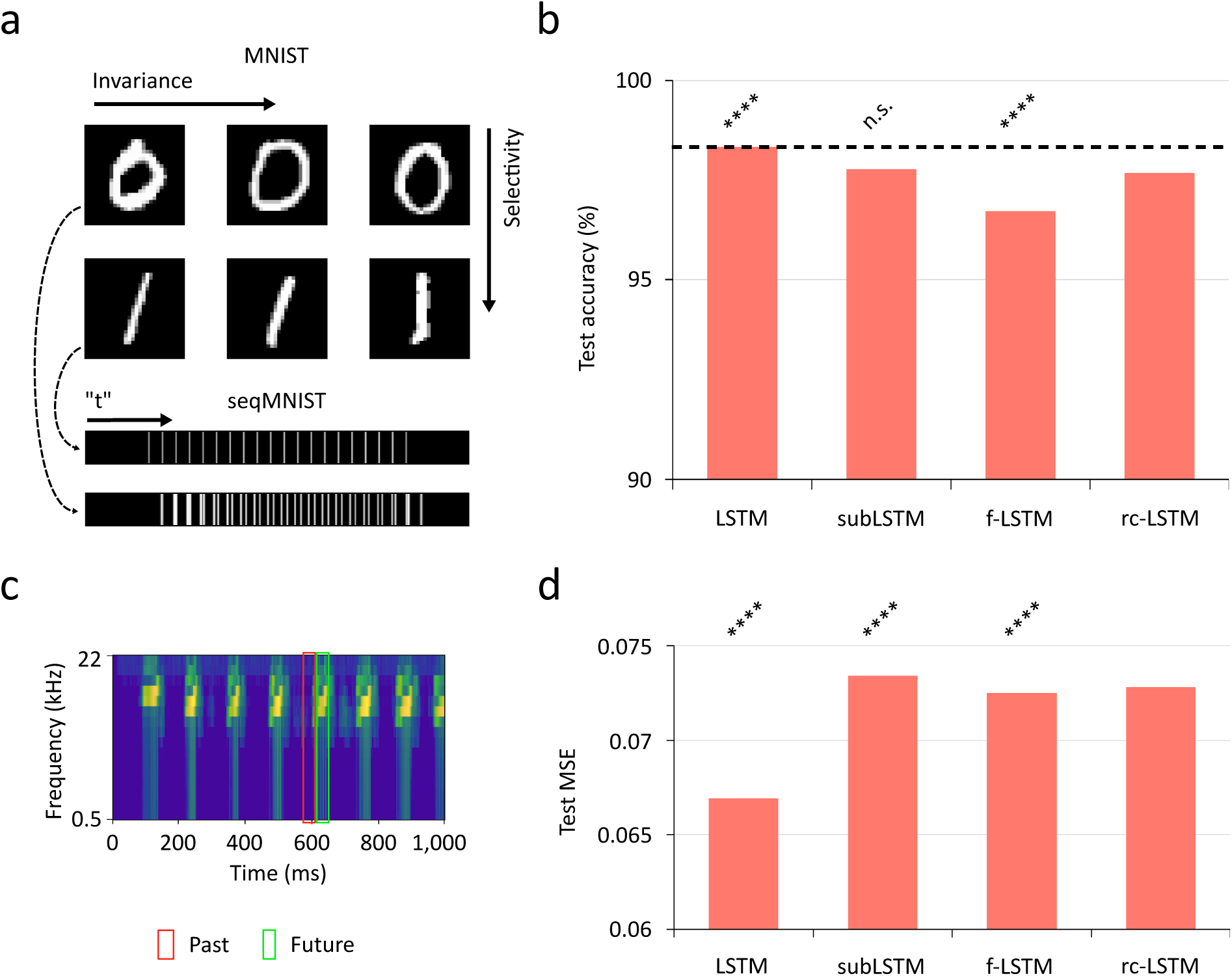
The f-LSTM and rc-LSTM models perform well on both a machine learning benchmark task and an auditory sequential task. (a) Illustration of the MNIST sequential classification task. 28x28 pixel images representing handwritten numerical digits 0-9 were transformed to 1D vectors and fed as sequences to the models. The goal is to train the models to distinguish the digits while maintaining invariance across different handwritten implementations. (b) Model accuracy on the task. All models perform similarly well, with only slight differences in accuracy. Asterisks indicate a significant difference between each of the models and the rc-LSTM (McNemar’s test; n_10_=141, n_01_=75; n_10_=136, n_01_=127; n_10_=113, n_01_=209; in comparison with the LSTM, subLSTM, and f-LSTM, respectively). (c) Illustration of the auditory temporal prediction task. Models are trained to predict a 20 ms cochleagram window into the future at each time bin. (d) Model mean squared error (MSE). The LSTM model is the best predictor of the future of the stimulus, followed by the f-LSTM model. Asterisks indicate a significant difference (Wilcoxon signed-rank test, n=58) between each of the models and the rc-LSTM.

However, most tasks performed by the auditory cortex will be substantially more complex and high-dimensional than the MNIST task. Hence, we tested our two new architectures on a sequential task more relevant to the auditory brain. Evidence suggests that the auditory cortex is optimized for the prediction of future input (temporal prediction)^37^. We trained models to predict the future of cochleagrams from their past, and found that the f-LSTM and the rc-LSTM models are better than the subLSTM model (Figs. 7c,d). As in the sequential MNIST task, the LSTM model performs best. This suggests that incorporating the biologically motivated subtractive gating worsens performance on a complex task, but that adding further biological features may somewhat compensate.

## Discussion

We developed a new gated recurrent model, the f-LSTM model, employing corticofugal-like input gating, to model auditory cortex responses to natural sounds. This architecture outperformed all other models in predicting the responses to natural sounds of units recorded in primary (A1) and secondary (PEG) cortex of awake ferrets. Having observed a larger benefit for the f-LSTM model in deep, corticofugally-projecting layers of A1, and inspired by the high neuronal density in superficial layers, we further improved A1 prediction performance with the rc-LSTM model, developed from the f-LSTM model by adding an expansion layer. Moreover, using the f-LSTM model, we quantified the use of memory in A1 and PEG, finding substantially longer temporal integration windows in the latter. We also identified two prominent natural sound features evoking responses that are better captured by models with than without memory: abrupt changes in sound level and long periods of quiet or silence. Finally, we evaluated our new models on two sequential tasks, finding comparable performance with established architectures.

Auditory models built on biological features can be used to ascertain which aspects of the anatomy drive successful neural response predictions, and therefore contribute to the processing of natural sounds. Given the success of our f-LSTM and rc-LSTM models, we hypothesize that the circuit involves different stages of the auditory pathway and layers of A1, including corticofugal-mediated gating, dimensionality expansion, and a gated memory system (Fig. 8).

**Figure 8.**
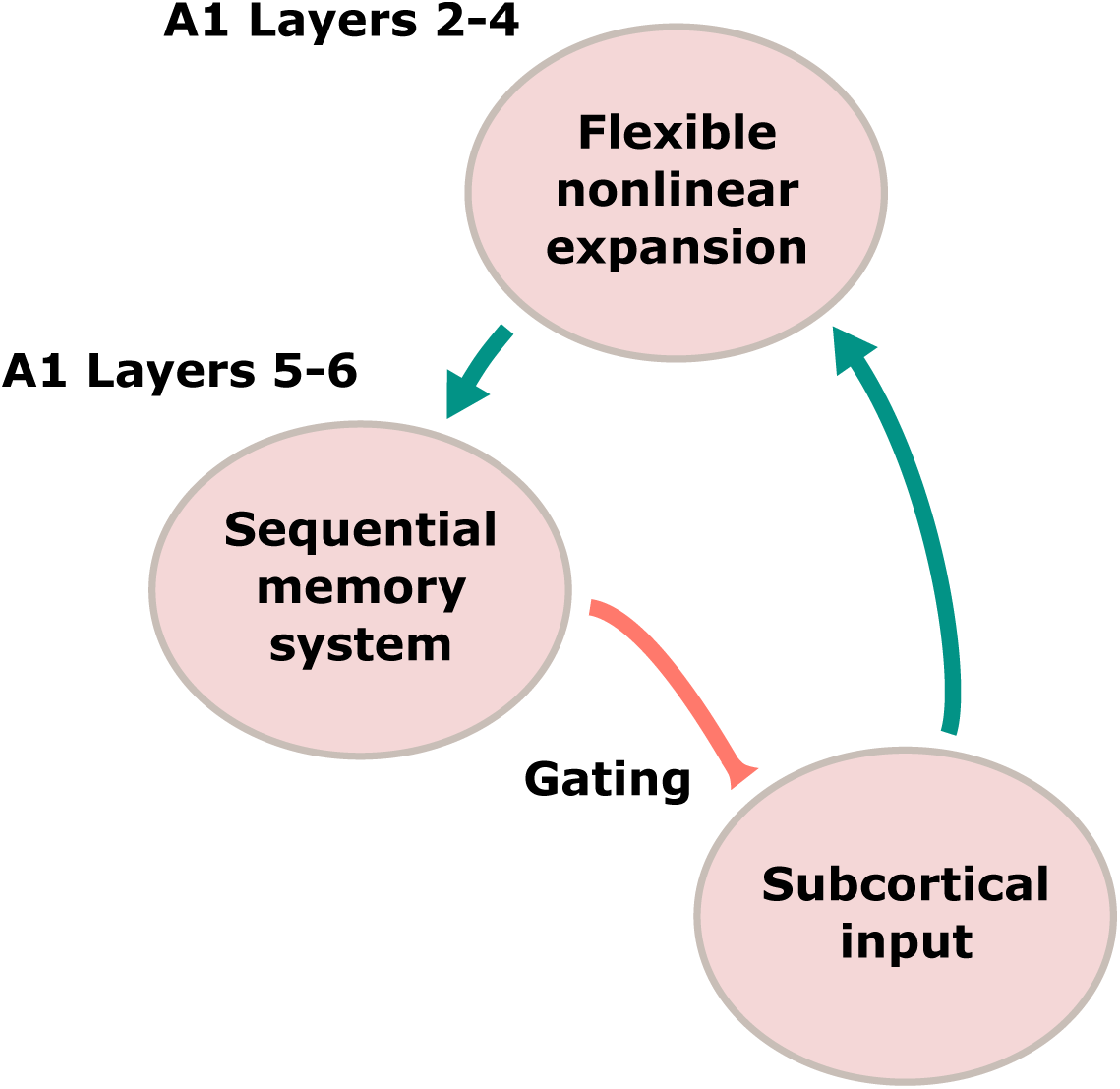
A proposed functional circuit for auditory processing. Subcortical input from the brainstem arrives via the thalamus at A1 in L4 and L2/3, which act as a dimensionality expansion step. Neurons in these layers project to deep layers (L5, L6), which act as a gated memory system. Corticofugal projections then carry information acquired at different timescales back to the subcortical centers, exerting top-down regulation of sound processing.

In comparison to other models of auditory cortical responses to natural sounds in awake ferrets, our f-LSTM and rc-LSTM models produce the highest CC_norm_ values seen in the literature for A1 and PEG. The convolutional models of Pennington and David (2023) come close, but rely on unrealistically long delay lines. Of the few auditory modelling studies incorporating memory-like mechanisms^7,13,15,17,38,39^, only two used explicit gated recurrency to predict extra-cellular spiking responses. Ahmed et al. (2025) found that a network with LSTM layers was the best predictor, amongst several speech recognition networks, of squirrel monkey responses to speech and primate vocalizations ^39^. Rançon et al. (2025) fitted GRU and LSTM models to the NAT4-A1 and NAT4-PEG datasets (among others), finding an advantage compared to feedforward models but obtaining lower CC_norm_ values than Pennington and David (2023) ^7^. Our results suggest that pairing gated memory with biological features - subtractive gating, corticofugal-like feedback, and superficial-layer expansion - enables better prediction performance than standard gated recurrency or convolutional models.

Subtractive gating has been linked to excitation-balancing inhibition within both canonical microcircuits in the cortex ^8^ and macrocircuits regulating working memory across whole brain regions ^40^. Other mechanisms that support memory, such as short-term plasticity^41^ and synaptic depression ^42^, have previously been incorporated into auditory cortical models, improving their prediction performance ^13,17^. Modelling work suggests that these mechanisms could be the consequence of inhibitory gating at the dendritic level ^43^. Further work is therefore necessary to determine whether gated models are faithful to the biology of the auditory cortex, or if they are simply capturing other memory mechanisms.

Corticofugal projections have been shown to influence sound processing along the auditory pathway^22–26,44–46^, including in processes specifically involving memory, like meta-adaptation to repeated changes in sound statistics ^45^ and repetition enhancement in the midbrain ^46^ and thalamus ^25^. A model combining memory with corticofugal-like top-down control can potentially capture these processes.

Superficial-layer expansion may bear some relation to the kernel trick, an efficient way to project data into higher dimensions, which is used in machine learning to process data that is not linearly separable using linear algorithms ^47^. High-dimensional representations subserving processing similar to the kernel trick have been reported in the visual cortex ^48^ and hypothesized in the auditory cortex^49^. This ability may (at least partially) underlie the success of the rc-LSTM model. A corresponding L3 - L5 circuit in the auditory cortex of mice (part of a wider L3- L5 - IC pathway that also involves corticofugal projections) has been implicated in behavioural inhibition of the acoustic startle response ^50^, which involves a form of sensory memory.

We found two stimulus features that evoke responses that are better captured by models with memory. Abrupt changes in sound level, which include sound onsets and offsets, have been implied in various species in sound duration encoding, perceptual grouping, and behavioral saliency^51,52^. Sensitivity to the duration of quiet or silent periods is, to our knowledge, a new phenomenon. Neural activity ramping up during periods of silence in sounds (up to 300 ms) has been previously reported in the auditory cortex of rats during an interval discrimination task^53^, but not as far as we aware in passively listening animals. This activity could reflect anticipation mechanisms ^53^ for certain upcoming behaviorally relevant sounds, such as the vocalizations of fellow ferrets.

The auditory cortex also shows numerous other memory phenomena, such as stimulus-specific adaptation ^3,54^, stimulus-specific facilitation ^25,55^, temporal context dependence at several timescales ^4,5,56^, and auditory working memory^57^. These processes, spanning timescales of tens of milliseconds to minutes, may all rely on some form of gating, and can be captured in principle by our models.

As in A1, the f-LSTM model is also the best predictor of PEG responses. However, in NAT4-PEG, the dataset obtained from this region of secondary auditory cortex, we did not find a correspondence between performance and neuronal depth, and the rc-LSTM performs worse than the f-LSTM model. In mice, secondary auditory cortex shows a less clear laminar organization than primary cortex, which is also more granular^58^. Our proposed circuit may therefore be more likely to exist in A1.

We also found longer memory retention in PEG than in A1. This confirms recent evidence from ferret recordings showing longer temporal integration windows in PEG compared to A1^59^. PEG is thought to be an intermediate stage between A1 and dorsolateral frontal cortex (dlFC) where acoustic features combine to form abstract representations of sound meaning ^60^. This involves integrating information at various timescales, and is likely supported by long memory retention.

Testing models on the sequential MNIST benchmark task, we find that the more biologically realistic models approach the performance of the LSTM model, but they still do not match it. Networks of LSTM units have for many years been the state-of-the-art for processing sequential data in machine learning and natural language processing. Therefore, our results suggest that the biological constraints imposed on our models may limit their capacity on a machine learning benchmark such as the sequential MNIST classification task. However, for the more complex high-dimensional temporal prediction task, which is more closely related to operations performed by the brain, the corticofugal-like feedback compensates somewhat for the constraints of the subLSTM model, enabling it to move towards the better performance of the LSTM model. This suggests that some of the complexity of the auditory system may be to do with compensating biological limitations.

While our new models improve upon all previously published alternatives, they have some limitations. First, they bypass all stages of the auditory pathway between the auditory nerve and the cortex. Incorporating hierarchical features based on subcortical responses has been shown to improve the prediction accuracy of cortical units in auditory models^38,61^. PEG, as a secondary area, would likely benefit even more from adding such hierarchical steps to the models, as suggested by recent work using A1 representations of stimuli to predict PEG responses^62^. Thus, stacked neural networks with a stage representing A1 may be more appropriate to model PEG. Exploring more biologically detailed cochleagrams ^18^ may also be useful. The rc-LSTM model and its associated circuit in Fig. 8 could also account more for differences between layers. The memory region corresponds to L5 and L6 pooled together, but these operate over different timescales ^63^. Indeed, we find a clear advantage of the f-LSTM model at a depth around L5 but not L6. In addition, L2, L3, and L4 are considered as one expansion stage in the circuit, overlooking the differences in connections between them ^64,65^.

We developed new models predicting ferret auditory cortical responses to natural sounds, that improve upon all previously published alternatives. Our models incorporate memory and, in the case of our best performing models, biological detail in the form of inhibitory-like gating, corticofugal projections, and dimensionality expansion, suggesting that these features are all involved in the functional circuits responsible for natural sound processing. Modelling these circuits may help to pinpoint causes and solutions for auditory disorders and improve speech processing algorithms.

## Methods

### Datasets

We fitted models to three previously published datasets of auditory cortical responses in awake ferrets (Table 1). The NAT4-A1 and NAT4-PEG datasets were recorded in response to a large set of 1-second long natural sounds^16^. As their names suggest, they were recorded in primary auditory cortex (A1 and anterior auditory field) and secondary auditory cortex (PEG), respectively. For convenience, we refer to the former as A1. The NS2 dataset was recorded in A1 in response to a smaller set of 4-second long natural sounds^17^. NAT4-A1 and NAT4-PEG contain both single- and multi-unit responses, whereas NS2 only contains single-unit responses. The stimuli used include animal sounds, human speech, music, and general and lab environment sounds.

**Table 1.** Dataset information.

| Dataset | Stimuli<sup>1</sup> | Repeats<sup>1</sup> | Duration (s) | Publication |
| --- | --- | --- | --- | --- |
| NAT4-A1 | 573/18 | 1/20 | 1 | Pennington and David (2023) |
| NAT4-PEG | 573/18 | 1/20 | 1 | Pennington and David (2023) |
| NS2 | 281/18 | 1/10 | 4 | Lopez-Espejo et al. (2019) |

<sup>1</sup> Most stimuli were presented to the ferrets only once, and the rest were presented over multiple (10 or 20) repeats

For all three datasets, stimuli were presented to ferrets via free-field speakers at a sampling rate of 44.1 kHz. Extracellular spiking activity was recorded with silicon multielectrode arrays (UCLA probes) for the two NAT4 datasets, and with tungsten microelectrodes (FHC) for the NS2 dataset. More detail about each dataset can be found in their respective linked repositories (https://zenodo.org/records/7796574, https://zenodo.org/records/3445557) and associated publications ^16,17^.

### Pre-processing

For all datasets, stimuli were available as sound pressure waves, and responses as spike times. We converted spike times to peri-stimulus time histograms (PSTHs), binned at 4ms time resolution. PSTHs were averaged over repeats if needed.

We converted sound pressure waves to time-frequency *cochleagrams*, spectrogram-based representations designed to approximate the frequency decomposition process taking place in the cochlea (Supplementary Fig. S1). The cochleagram transformation we used (*spec-power* ) has been shown to enable a good level of fitting performance across several encoding models of auditory cortex^18^. We binned the cochleagrams at 4 ms (like the PSTHs), and used 8 or 16 frequency channels (see Supplementary Information) spaced logarithmically between 500 and 22,000 Hz.

For each dataset, we normalized cochleagrams to have zero mean and unit variance over the whole training set. All models explored in this study produce a prediction of neural activity at each time bin and are fed a spectrotemporal window of stimulus history preceding the time bin, spanning all frequencies and a range of latencies (delay lines). We zero-padded sound pressure waves to their left with a span corresponding to the window.

We also calculated a *noise ratio* (NR) statistic to evaluate the variability of each recorded unit’s response over multiple repeats of the same stimuli ^66^. We selected a noise ratio of 40 as the threshold above which units were excluded from the fits. We also excluded units if they fired, on average, *<* 0.5 spikes/s over all multi-repeat stimuli. Finally, to eliminate inconsistent units, we excluded them if the variance of their mean firing rate over all repeats exceeded a threshold (10*^−^*^4^, chosen based on mean variance) for more than half of the multi-repeat stimuli. The final numbers of units that survived all exclusion criteria were 383, 195, and 180 in the NAT4-A1, NAT4-PEG, and NS2 datasets, respectively.

### Models

We used 12 models in this study, all of which were implemented as population neural network architectures, which estimate the responses of all units in a dataset at once^16^. The feedforward models have all been previously published, and they are briefly explained in the Supplementary Information.

All recurrent and gated recurrent models, except the f-LSTM and the rc-LSTM, have the same structure: a layer of recurrent or gated recurrent units (Supplementary Fig. S3) followed by a linear mapping with a sigmoid nonlinear activation function to obtain the final PSTH predictions (Supplementary Fig. S2). All network units also have an associated bias parameter.

The RNN model uses standard Elman recurrent units ^28^. The MGU, GRU, LSTM, and subLSTM models were named after the type of gated recurrent units they use. Gates consist of sigmoidal functions driven by the units’ input and memory state. The outputs of the gates are then multiplied with the input and memory state themselves. This enables only relevant information to be selectively input and retained in memory, dependent on current input and memory.

The Elman recurrent unit does not use any gating. Its input (*x_t_*) and recurrent memory state (the hidden state, *h_t_*) are passed through a tanh function to obtain the next hidden state^28^. In contrast, the LSTM unit^19^ contains three gates (input, forget, output) and has two recurrent memory states: the hidden state (*h_t_*) and the cell state (*c_t_*), which contains the *long* memory referred to in the name. The gates operate on the memory states and on incoming inputs (*x_t_*), allowing the latter to add to the memory or resetting the cell state to a default, for example. The GRU is a simplified version of the LSTM unit^20^. Its two gates (update, reset) fulfill the same functions as the three LSTM gates, and the hidden state (*h_t_*) carries all the memory (there is no cell state). The MGU is a further simplification of the LSTM unit and the GRU^21^. It only contains one gate (forget) and has one memory state (*h_t_*).

The subLSTM unit (Fig. 9) is a biologically inspired alternative to the LSTM unit^8^. It contains the same three gates, but two of them (input, output) operate subtractively, rather than multiplicatively, on the memory and the incoming input. Subtractive gating approximates the action of inhibitory neurons in the cortex better than multiplicative gating^8,67^. The subLSTM unit is described by the following equations:

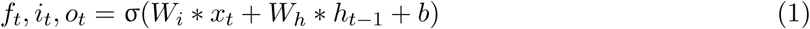

**Figure 9.**
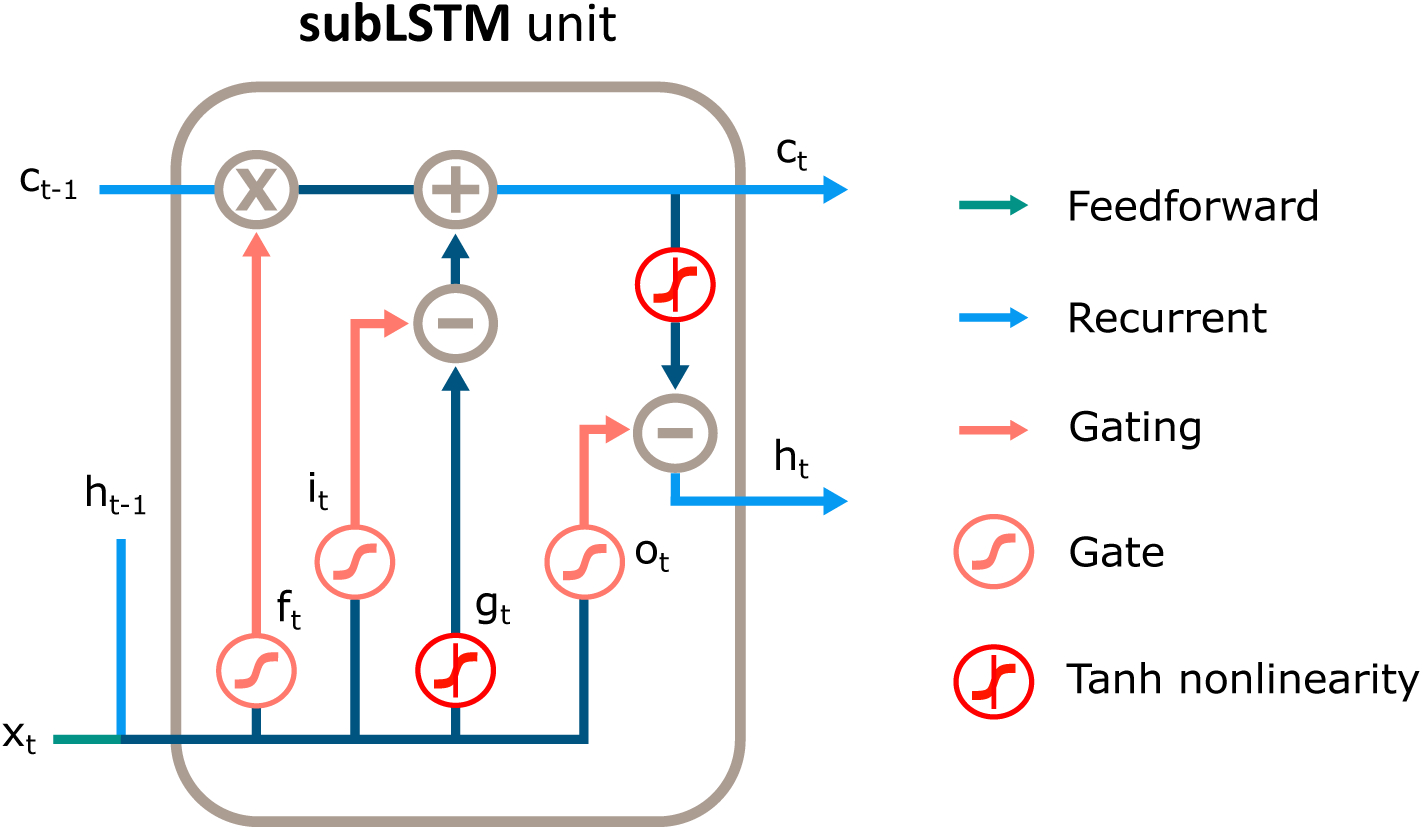
Schematic of the subLSTM unit ^8^. It receives an input *x_t_* from the preceding layer in the network and passes two memory states (*h_t_*, hidden state and *c_t_*, cell state) across time steps. It has three gates: forget (*f_t_*), input (*i_t_*), and output (*o_t_*); *g_t_*is the candidate cell state.

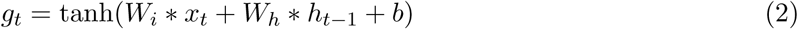

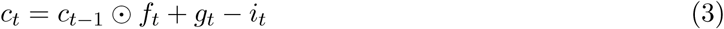

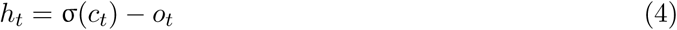

where *x_t_* is the input to the unit from the previous layer, *h_t_* is the hidden state, and *c_t_* is the cell state; *f_t_*, *i_t_*, and *o_t_* are the forget, input, and output gates, respectively, and *g_t_* is the candidate cell state. *W_i_*and *W_h_*are the weights carrying the new input and the existing hidden state, respectively. These are different for each gate and for the candidate cell state. The associated biases are also different for each of these, but are indicated by *b* for simplicity. In all the equations that follow, *b* indicates bias parameters. The operators ⊙ and σ() indicate element-wise multiplication and the sigmoidal function, respectively.

To obtain the final firing rate predictions, the hidden state is passed through a linear mapping and a sigmoidal activation as follows:

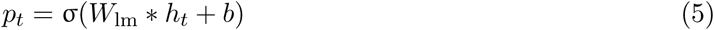

where *p_t_*is the predicted firing rate and *W*_lm_ are the linear mapping weights.

The f-LSTM (Fig. 1b) and rc-LSTM (Fig. 4e) models also have a hidden layer of subLSTM units followed by a linear mapping and a sigmoidal activation at the output. However, in the f-LSTM model, the hidden state of the subLSTM units and the input cochleagram are fed into an additional gate, also implemented as a sigmoidal function. This *input gating* mechanism allows only relevant portions of the cochleagram to be fed to the gated recurrent layer. It is implemented as follows:

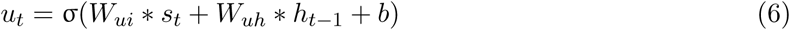

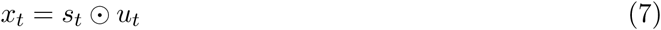

where *u_t_*is the input gate, *s_t_*is the input stimulus, and *W_ui_*and *W_uh_* are the weights to the input gate from the input and the hidden state, respectively.

The rc-LSTM model also uses input gating, but the gated cochleagram is passed through a linear *expansion* layer and a sigmoidal nonlinearity prior to the subLSTM one, as follows:

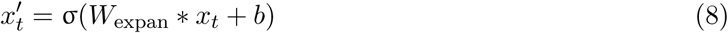

where *W*_expan_ are the expansion layer weights, *x_t_*is the gated input (equation 7) and 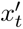 is the expanded input fed to the subLSTM layer.

### Modelling pipeline

#### Model fitting procedures

We initialized all model parameters (weights and biases) to their PyTorch default, which samples parameter values for each layer from a uniform distribution between 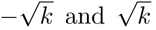, where 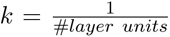. The only exception is the recurrent weight matrix of the RNN model, which we initialized to an identity matrix following a popular method ^36^. We also initialized the memory states of recurrent and gated recurrent models to 0. We fitted parameters with the NAdam optimization algorithm for stochastic gradient descent^68^ with a mean squared error (MSE) loss function ^69^, calculated between model output (predicted PSTH) and target (actual PSTH), whose gradients we obtained via backpropagation^70^. In addition, we trained all models with gradient clipping (with a threshold at 1) to prevent the exploding gradient problem ^71^. The values of the training hyperparameters are reported in the Supplementary Information.

During training, we fed the models each stimulus-response snippet separately, randomizing their order at each epoch. At the end of each snippet, we erased the memory of recurrent and gated recurrent models. To prevent overfitting to training data, we used L1-norm regularization on all models ^72^, trialing several regularization strength (*λ*) values (see Supplementary Information).

### Training, validation, and testing datasets

To explore the space of hyperparameters and evaluate model prediction performance in an unbiased manner, we split each dataset into a training set, a validation set, and a test set ^73^. In NAT4-A1 and NAT4-PEG, we used 580 stimulus-response snippets for training, 7 for validation and testing, and 4 just for testing. All validation and testing snippets were from multi-repeat stimuli, as were 7 out of the 580 training snippets (the rest corresponding to single-repeat stimuli). In NS2, we used 288 snippets for training (again, 7 of which from multi-repeat stimuli), 7 for validation and testing, and 4 just for testing. The test set was held out from any fitting or tuning procedure, and was solely used in the evaluation of the final model fits. We used the training and validation sets to determine the *λ* (regularization strength) hyperparameter, choosing the fitted parameters associated with the *λ* value that maximized prediction performance on the validation set. Finally, we evaluated models on the held out test set combined with the validation set.

The training and validation sets were also used to determine the number of training epochs, the number of hidden layer units, the number of frequency channels, and the length of delay lines (all detailed in the Supplementary Information).

### Model performance measure: the normalized correlation coefficient

We used the normalized correlation coefficient (CC_norm_) to measure model performance ^29,30^. This takes the raw correlation coefficient (CC_raw_) between the actual PSTH of a recorded unit and the model’s prediction of it, and normalizes it by the maximum achievable correlation coefficient (CC_max_) for that unit given how variable its responses are over multiple repeats of the same stimuli. The CC_norm_ is 0 when the model prediction is uncorrelated with the actual neural response, and 1 when it is as correlated as possible given this variability. By definition, this calculation is only possible on multi-repeat stimulus-response snippets. For this reason, we constructed PSTHs of multi-repeat responses used to compute CC_norm_ separately for each repeat. For more detail on how the measure is calculated in practice, see Schoppe et al. (2016).

### Model analysis

#### Statistical methods

For paired statistical comparisons of prediction performance between models (Figs. 2c, 3c, 4f,g, and 5c), we used non-parametric two-tailed Wilcoxon signed-rank tests. We also used these tests for paired comparisons of model performance on the temporal prediction task (Fig. 7d).

For comparing model performance in different cortical layers, we used an unpaired two-tailed t-test to assess whether improvements in performance between the f-LSTM and the 2D-CNN models were different in L5/6 compared to L2/3 and L4 (Figs. 4a,b).

To assess the significance of linear correlation between the performance differences between models and the abrupt changes and quietness duration measures (Figs. 6d-g), we used an unpaired two-tailed t-test as the built-in statistical test of the ordinary least-squares regression.

To compare model performance (accuracy) on the sequential MNIST classification task (Fig. 7b), we used paired two-tailed McNemar’s tests.

Statistical significance for all tests was set at p*<*0.05. When used, asterisks indicate the level of significance as follows: * indicates p*<*0.05, ** indicates p*<*10*^−^*^2^, *** indicates p*<*10*^−^*^3^, and **** indicates p*<*10*^−^*^4^ and below.

### Depth analysis

We obtained the values of depth within cortex for a subset of units in each dataset from Stephen David. To relate performance differences between models to the recording depth, we took the mean of the CC_norm_ and depth of each unit over 200 µm-wide windows of depth, overlapping by 50 µm. We chose these values to have clear comparisons between models while preserving good resolution of depth across cortical layers.

The distribution of depths for units in the NAT4-A1 dataset was wider than for the NS2 dataset (Fig. 4c). To maintain comparability across datasets, we excluded the non-overlapping units in NAT4-A1 from the analysis.

### Temporal integration analysis

To analyze and compare temporal integration properties of A1 and PEG, we used the NAT4-A1 and NAT4-PEG datasets, as they contained the same set of stimuli. We first split each stimulus-response snippet into equal-length non-overlapping segments, or ”chunks”, of either 1/2, 1/3, 1/4, 1/8, 1/9, 1/16, 1/27, or 1/32 (also 1/64 and 1/128 for the NS2 dataset) of the length of each snippet. We then fitted the f-LSTM model separately for each chunk length, applying the same training, validation, and testing procedure used in the main model fits. We kept the same hyperparameters as in the original f-LSTM fit, except for the regularization strength (*λ*), which we optimized for each chunk length.

We fitted the relationship between model performance and chunk length with a power-law function, of the form:

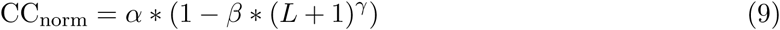

where *L* is the chunk length in ms, and *α*, *β*, *γ* are the function’s parameters. We fitted the power law with a nonlinear least-squares algorithm. The *α* and *β* parameters were bounded between 0 and 1, and the *γ* parameter was bounded between -1 and 0. We were limited by the length of the stimuli to a maximum chunk length of 1 second, but auditory cortical context dependence has been previously shown to span longer timescales^4^. In addition, most data points corresponded to short (*<* 200 ms) chunk lengths. To avoid biasing the fits to these points and obtain more meaningful predictions of neuronal behaviour over long timescales, we used the *DenseWeight* algorithm, which assigns larger weighting to residuals of rare data points in unbalanced datasets ^74^.

We calculated the chunk length at which model performance reaches 80% of its maximum as:

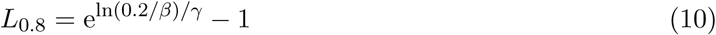

### Stimulus features analysis

To investigate reasons why the f-LSTM model improves upon its feedforward counterparts, we first examined the most improved units qualitatively, and identified two activity patterns: responses to abrupt changes and sensitivity to the duration of quiet periods in sounds.

To quantify the prevalence of abrupt changes and the duration of quiet periods in each stimulus, we developed two measures. For the abrupt changes measure, we first computed a high-pass filtered version of each stimulus cochleagram, applying 4^th^ order Butterworth high-pass filters to each frequency band. Filters had a cutoff frequency (*f*_c_) that depended on the band’s center frequency (*f* ) as follows ^38,75^:

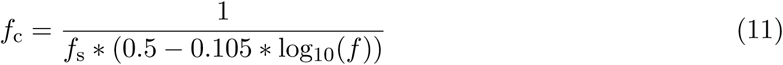

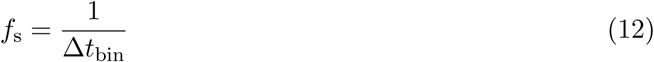

where *f*_s_ is the cochleagram ”sampling frequency”, calculated as the inverse of the time bin size Δ*t*_bin_ in seconds. We then positively half-wave rectified the high-pass filtered cochleagrams, and computed the abrupt changes measure (*A*) as follows:

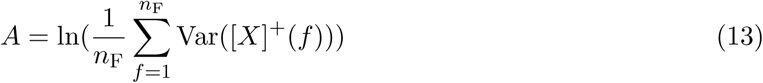

where *n*_F_ is the number of frequency channels, [*X*]^+^ is the half-wave rectified cochleagram, and Var() is the variance operator.

To obtain the quietness duration measure for each stimulus, we first z-scored each frequency channel in the cochleagram. We then built a quietness counter for each channel by counting the number of time bins with negative amplitude since the last positive-amplitude bin. At each bin with positive amplitude, the counter was reset to zero. Putting together the quietness counter for each frequency channel, we obtained a quietness counter matrix, *M* , from which we computed the quietness duration measure as follows:

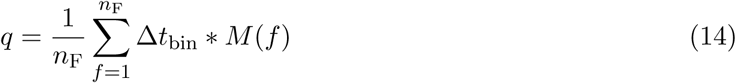

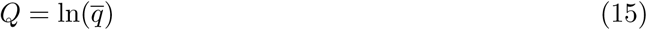

where *Q* is the quietness duration measure in ms and *q* is the mean quietness counter across frequency channels.

We then calculated the correlation coefficient (CC_raw_) between the neural responses and the model predictions for the f-LSTM and for the best feedforward model (the 2D-CNN and 1D-CNN for NAT4-A1 and NAT4-PEG, respectively) for each neuron and stimulus. As we covered all stimuli in this analysis, including those only presented to the ferrets once, we could not use CC_norm_. For each stimulus, we took the mean of the CC_raw_ across units, and calculated the difference in the mean CC_raw_ between the f-LSTM and the feedforward model. We normalized this difference by its mean over all stimuli. We also sorted the stimuli by each metric, splitting the results in deciles and computed the means of the normalized CC_raw_ difference and of the measures for each decile.

We fitted the dependence of the normalized CC_raw_ difference on each measure with a linear function using ordinary least-squares (OLS) regression (Figs. 6d-g).

### Sequential MNIST benchmark task

We obtained the sequential MNIST benchmark task results using the methods detailed in Costa et al. (2017). A pipeline for fitting models to the sequential MNIST dataset was im-plemented here: https://github.com/neuralml/pytorch-subLSTM. We adapted the code for the f-LSTM and rc-LSTM models for use in the pipeline.

### Temporal prediction task

In the auditory temporal prediction task, the models were fed 32 ms of cochleagram history at each time bin, and had to predict the cochleagram 20 ms into the future. We used a large dataset of 572 natural sounds, converting them to cochleagrams with 8 frequency channels and binned at 4 ms. We normalized cochleagrams to have zero mean and unit variance over the whole training set. The models all had 150 hidden units. We fitted parameters with the NAdam optimization algorithm (10*^−^*^4^ learning rate) with an MSE loss function, with gradients calculated via backpropagation. Training lasted 250 epochs. We applied L1-regularization to prevent overfitting. We used 10% of cochleagrams for testing, and the remaining 90% were split 90-10 in 4 different ways for cross-validation. Cross-validation was used to determine the regularization strength (the values trialed were 1.28 × 10*^−^*^8^, 2.56 × 10*^−^*^9^, 5.12 × 10*^−^*^10^, 1.024 × 10*^−^*^10^, 2.048 × 10*^−^*^11^, and 4.096 × 10*^−^*^12^) and the number of cochleagram frequency channels. Those were then used in the final evaluation on the test set (Fig. 7d).

### Software used

We built, trained, and evaluated all models in Python 3.10, with the PyTorch library, using built-in objects from its neural network (torch.nn) sub-package^76^.

## Data availability

The datasets we used are available at https://zenodo.org/records/7796574^77^ (NAT4-A1 and NAT4-PEG) and https://zenodo.org/records/3445557^78^ (NS2). The depth information was provided to us by Stephen David. The MNIST dataset is publicly available online - we used the one from the TorchVision Python library. The natural sounds used in the temporal prediction task are publicly available and found on the Freesound (https://freesound.org/) website.

## Code availability

All the original code used to produce the results described in this paper can be accessed at: https://github.com/lm1017/corticofugal-LSTM. The original code for the sequential MNIST task, which we modified to include the f-LSTM and rc-LSTM models, can be found at https://github.com/miltonllera/pytorch-subLSTM.

## Acknowledgments

We would like to thank Kerry Walker and Rui Ponte Costa for helpful comments and feedback throughout this work. We would also like to thank Stephen David for providing additional information on the data used, and Milton Llera-Montero for help with the sequential MNIST classification task. A.J.K., B.D.B.W., and N.S.H. are supported by Wellcome Trust Principal Research Fellowship WT108369/Z/2015/Z. L.M. is supported by a University of Oxford Medical Sciences Graduate School Studentship (covered partly by the Department of Physiology, Anatomy and Genetics and partly by the UKRI Medical Research Council).

## Author contributions

Conceptualization: L.M., B.D.B.W, N.S.H., Model development: L.M., B.D.B.W., N.S.H.,

Model fitting: L.M., Model analysis: L.M., Writing: L.M., A.J.K., B.D.B.W., N.S.H., Supervision: A.J.K., B.D.B.W., N.S.H.

## Correspondence and materials requests

Those should be addressed to Lorenzo Mazzaschi.

