## Supplementary Information for "Corticofugal gated recurrency captures auditory cortical responses"

### Cochleagram transformation

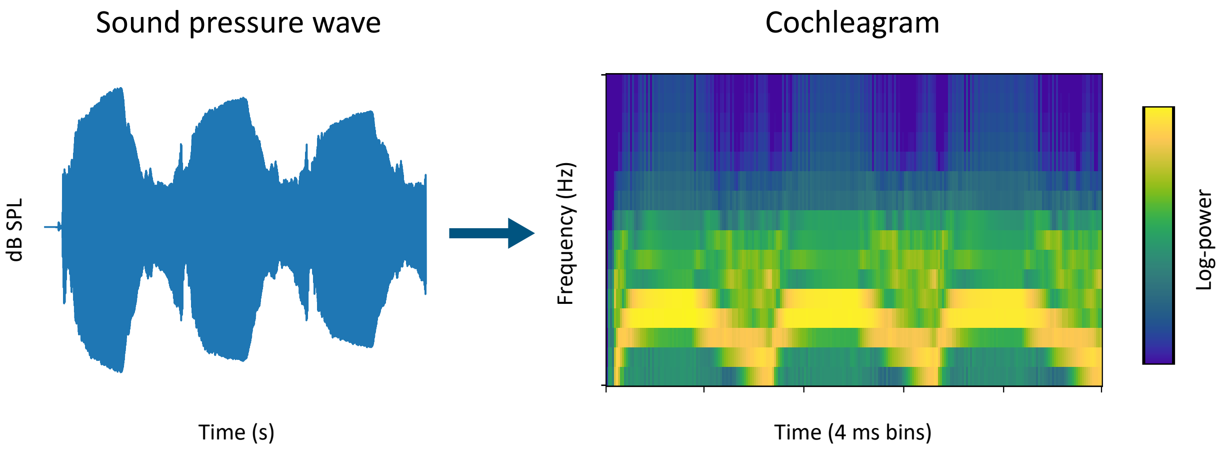


**Figure S1.** Sound wave and its corresponding cochleagram for one of the stimuli in NAT4-A1. This cochleagram has time bins of 4 ms and 16 frequency channels.

### Models

#### Feedforward models

The population linear-nonlinear (LN) model[16] was derived from the single-neuron LN model of Atencio et al. (2008). It consists of a neural network with one hidden layer, containing 1D convolutional filters convolved with the cochleagram along the time axis, with the result then summed over all frequency channels. A linear mapping relays the hidden layer activations to the output layer. To produce firing rate predictions, output unit activations are passed through a static sigmoidal nonlinearity. All units in the network have an associated bias parameter.

Our network receptive field (NRF) model is a population version of the NRF architecture developed by Harper et al. (2016). The NRF model is essentially a collection of single-neuron LN models arranged together in a one-hidden-layer, feedforward neural network. In contrast to the LN and the CNN models, which used convolutional filters, the NRF model was implemented with a linear hidden layer. Hidden layer activations are passed through static sigmoids before being pooled to the output layer via a linear mapping. As in the LN model, output unit activations are passed through static sigmoids to obtain the final predictions.

Several convolutional neural network (CNN) models were introduced by Pennington and David (2023). We implemented the 3 best architectures based on their performance in the original paper. These are the 1D-CNN, 1D-x2-CNN, and 2D-CNN models. The 1D-CNN model has two hidden layers: one convolutional layer with 100 1D filters spanning 248 ms, and one linear layer of 120 units. The 1D-x2-CNN model has two 1D convolutional layers, with 70 filters of 152 ms and 80 filters of 100 ms, respectively, and one linear layer of 100 units. Finally, the 2D-CNN model has 3 2D convolutional layers, each with 10 filters spanning 84 ms and 3 frequency channels, followed by a linear layer of 90 units. 2D filters are convolved with the cochleagram along the spectral and temporal axes simultaneously. In all three networks, the activations of each layer are passed through static sigmoid nonlinearities. Detailed schematics of these models can be found in the original publication[16].

#### Model and unit schematics

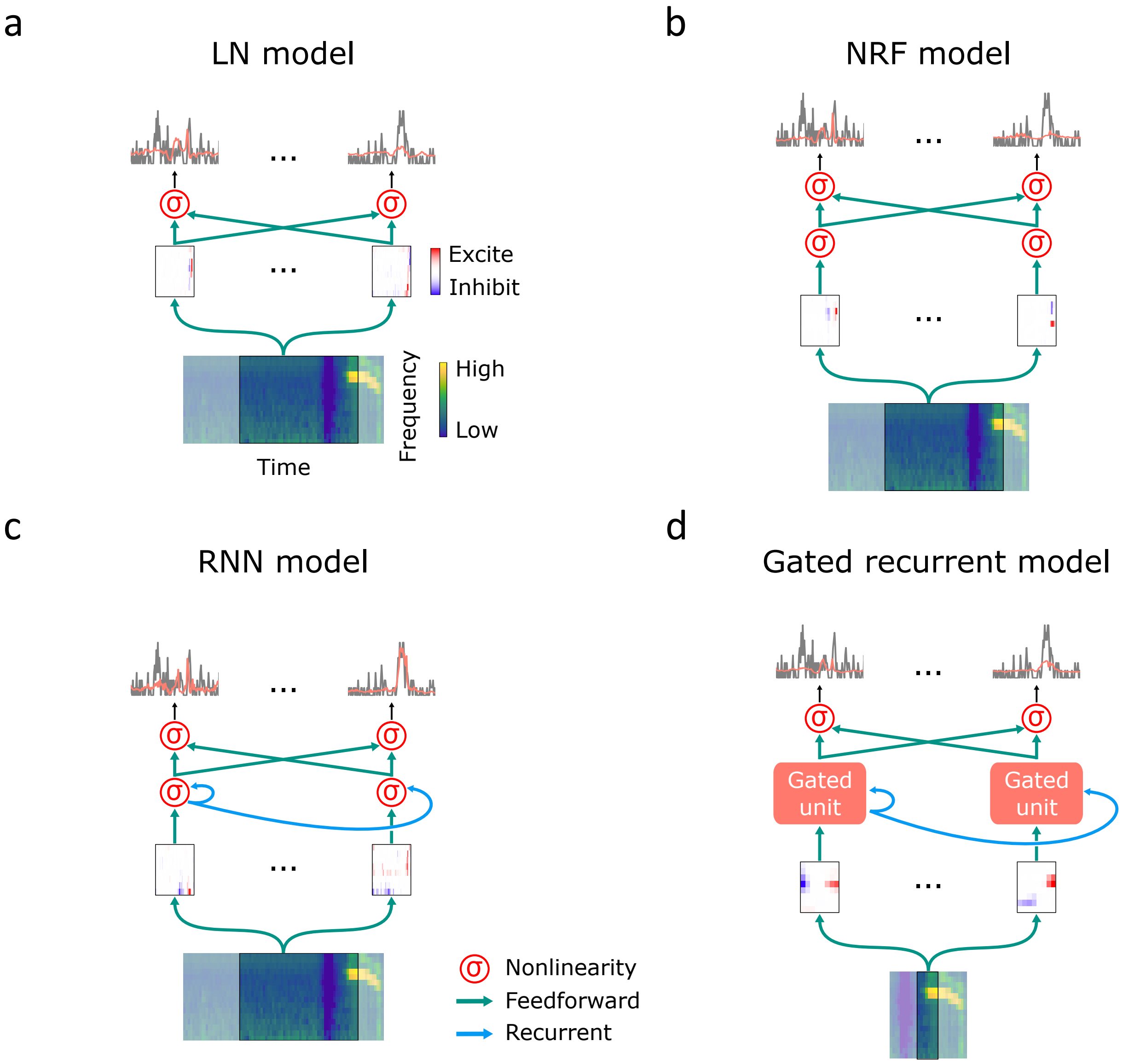


**Figure S2.** Schematics of the different architectures in order of complexity. LN model (a), NRF model (b), RNN model (c), and a general schematic of the gated recurrent models (d).

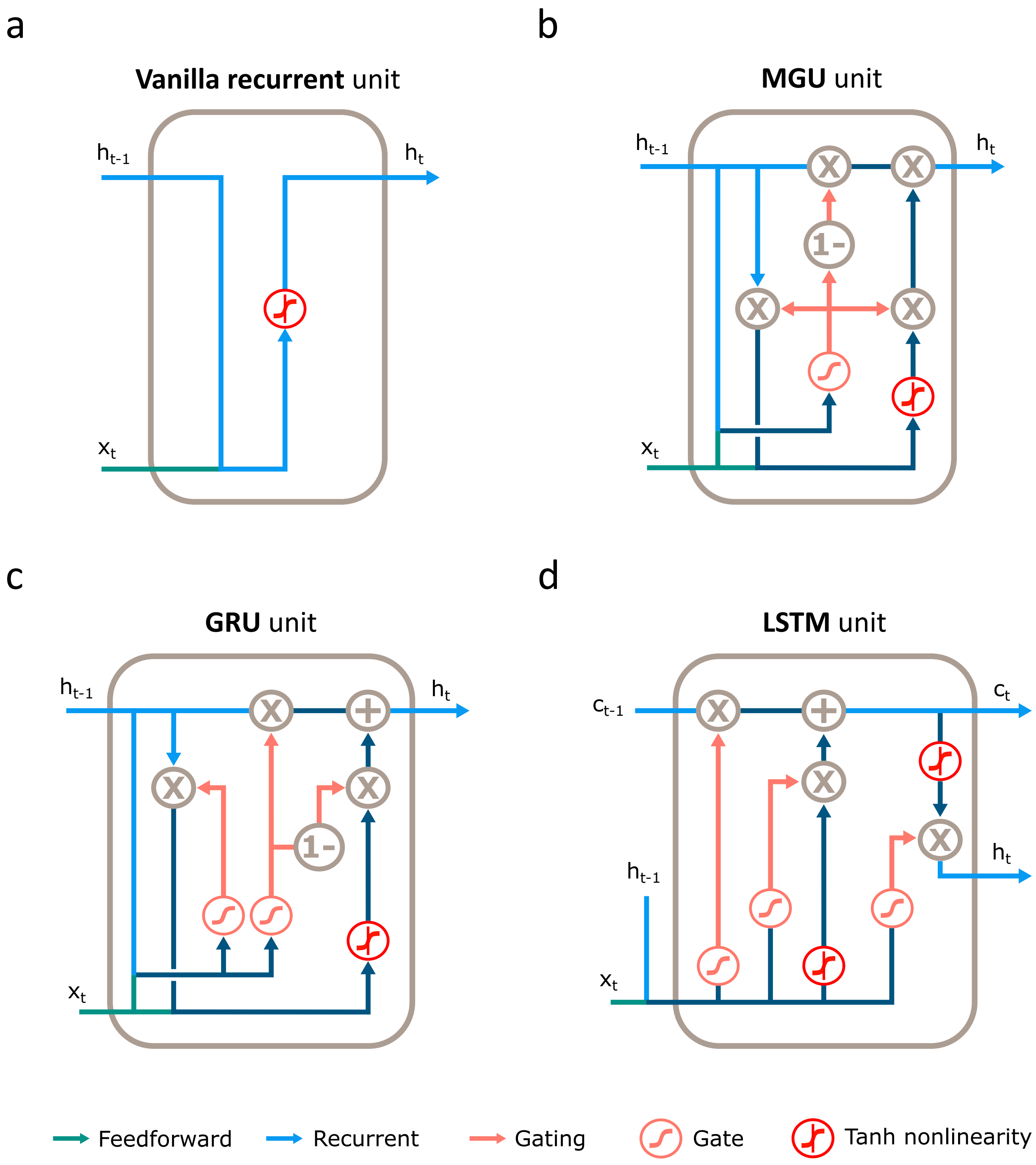


**Figure S3.** Schematics of recurrent and gated recurrent units used. (a) Vanilla recurrent unit. (b) Minimal Gated Unit (MGU). (c) Gated Recurrent Unit (GRU). (d) Long Short-Term Memory unit (LSTM). All units receive an input $x_t$ from the preceding layer in the network and pass a memory state $h_t$ (hidden state) across time steps. The LSTM unit passes an additional memory state $c_t$ (cell state).

### Example unit identifiers

The labels in the original datasets for the example units shown are: TNC020a-25-1 (Fig. 2a), TNC020a-16-1 (Fig. 2b), AMT018a-43-4 (Fig. 3a), AMT018a-28-3 (Fig. 3b), ARM019a-01-5 (Fig. 5a), ARM017a-39-4 (Fig. 5b), DRX008b-72-5 (Fig. 6a), and ARM022b-34-3 (Fig. 6b).

### Model performance and neuronal depth

When averaging depths and $CC_{norm}$ values over 200 μm depth windows, we ignored windows that included 3 or less units. As a result, a small number of units at each end of the depth range was excluded in Supplementary Figures S4a and S4c. This does not affect the plots in Figs. 4a and 4b.

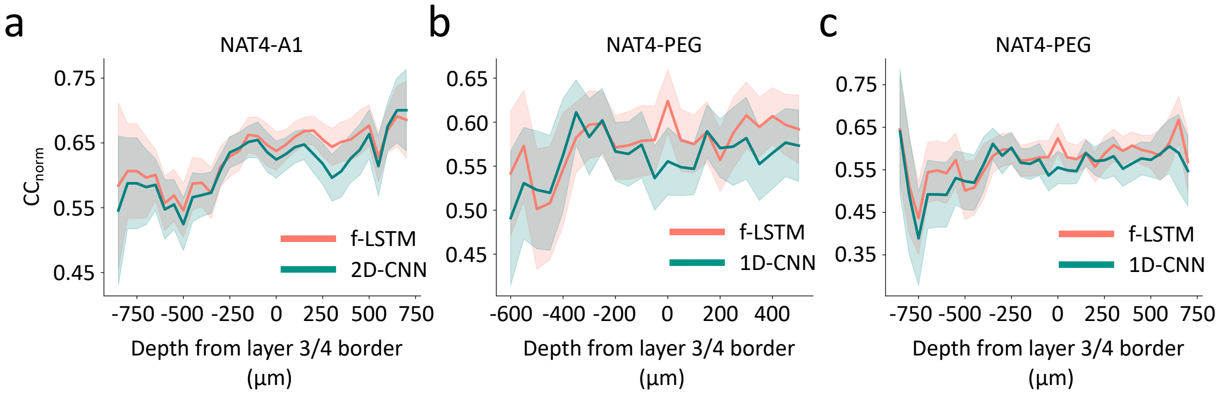


**Figure S4.** Model performance ($CC_{norm}$) plotted against neural depth. (a) Performance of the f-LSTM and 2D-CNN models against depth for the NAT4-A1 dataset, over the whole depth range available. (b) Performance of the f-LSTM and 1D-CNN models against depth for the NAT4-PEG dataset, over the range of depths of the NS2 dataset. (c) Same plot as in (b), but over the whole depth range available for NAT4-PEG. Negative numbers denote recording depths more superficial than the layer 3/4 border.

### Memory retention in the NS2 dataset

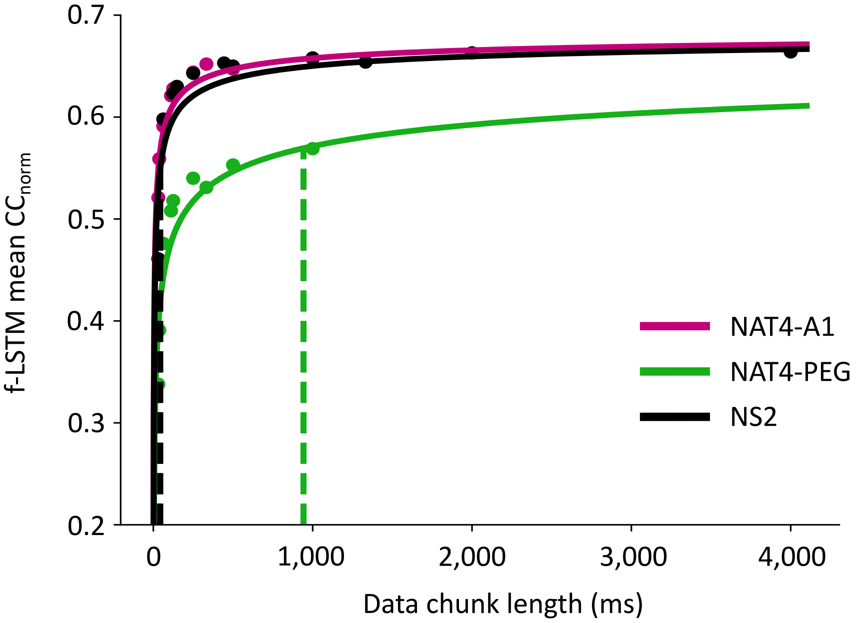


**Figure S5.** f-LSTM model mean $CC_{norm}$ plotted against chunk length, for all 3 datasets. The solid lines show power-law fits, whereas the dashed vertical lines indicate the chunk length at which the $CC_{norm}$ reaches 90% of its maximum value.

### Hyperparameter values

PSTHs and cochleagrams were binned in time at 4 ms. Cochleagrams had a varying number of frequency channels ($n_F$) depending on the model and dataset (Table S1). Cochleagram frequencies were always logarithmically spaced from $f_{min}$ = 500 Hz to $f_{max}$ = 22,000 Hz, where $f_{min}$ and $f_{max}$ are the minimum and maximum channel center frequency, respectively. We used different delay line lengths ($n_h$) for different models (Table S1). In models using convolutional filters, delay lines correspond to the time span of the filters; in non-convolutional models, delay lines correspond to the amount of history used in sound tensorization. All feedforward models have been previously published, and we used delay line lengths reported in the original articles after confirming their optimality with our fitting pipeline. For recurrent and gated recurrent models, we trialed several values and chose the one providing the best performance on the validation set (see Methods).

**Table S1.** Hyperparameters used in data pre-processing for each model and dataset

| Model | Delay lines (ms/$n_h$)[1] | $n_F$ | | |
|---|---|---|---|---|
| | | NAT4-A1 | NAT4-PEG | NS2 |
| LN | 200/50 | 16 | 16 | 16 |
| NRF | 200/50 | 8 | 8 | 8 |
| 1D-CNN | 248/62 | 8 | 8 | 8 |
| 1D-x2-CNN | 252/63 | 16 | 8 | 8 |
| 2D-CNN | 252/63 | 16 | 16 | 16 |
| RNN | 200/50 | 8 | 8 | 8 |
| MGU | 32/8 | 8 | 8 | 8 |
| GRU | 32/8 | 8 | 16 | 16 |
| LSTM | 32/8 | 8 | 8 | 8 |
| subLSTM | 32/8 | 8 | 8 | 8 |
| f-LSTM | 32/8 | 16 | 8 | 8 |
| rc-LSTM | 32/8 | 16 | 8 | 16 |

[1] Amount of stimulus history used by each model at any given time instant

We used different numbers of training epochs for each model and dataset (Table S2). We also trialed several values for the strength of L1-regularisation ($\lambda$): $1\times10^{-3}$, $2\times10^{-4}$, $1.17\times10^{-4}$, $6.84\times10^{-5}$, $4\times10^{-5}$, $2.34\times10^{-5}$, $1.37\times10^{-5}$, $8\times10^{-6}$, $4.68\times10^{-6}$, $2.74\times10^{-6}$, $1.6\times10^{-6}$, $9.36\times10^{-7}$, $5.41\times10^{-7}$, $3.2\times10^{-7}$, $6.4\times10^{-8}$, $1.28\times10^{-8}$, $2.56\times10^{-9}$, $5.12\times10^{-10}$. We kept the optimization learning rate fixed at $10^{-3}$ for all models and all datasets.

**Table S2.** Number of training epochs for each model and dataset.

| Model | Training epochs | | |
|---|---|---|---|
| | NAT4-A1 | NAT4-PEG | NS2 |
| LN | 100 | 100 | 150 |
| NRF | 100 | 100 | 1000 |
| 1D-CNN | 100 | 100 | 1000 |
| 1D-x2-CNN | 100 | 150 | 1000 |
| 2D-CNN | 150 | 150 | 1000 |
| RNN | 150 | 100 | 150 |
| MGU | 150 | 150 | 150 |
| GRU | 150 | 150 | 150 |
| LSTM | 150 | 150 | 150 |
| subLSTM | 150 | 150 | 150 |
| f-LSTM | 100 | 150 | 150 |
| rc-LSTM | 100 | 100 | 100 |

All models had a hidden layer of 150 units, except the 1D-CNN, 1D-x2-CNN, and 2D-CNN models, which were unchanged from their original implementation by Pennington and David (2023), with the length of convolutional filters adjusted to a time bin size of 4 ms (from 10 ms in the original publication). All convolution-based models had $n_\mathrm{F}$ as their input size; all others had $n_\mathrm{F} \times \mathrm{n_h}$ (due to cochleagram flattening, see Methods). Finally, all models had output layers with the number of units corresponding to the number of recorded units in the dataset being fitted.
